# Do African savanna elephants (*Loxodonta africana*) eat crops because they crave micronutrients?

**DOI:** 10.1101/673392

**Authors:** Susanne Marieke Vogel, Willem Frederik de Boer, Moses Masake, Anna Catherine Songhurst, Graham McCulloch, Amanda Stronza, Michelle Deborah Henley, Tim Coulson

## Abstract

1. Elephants can cause negative consequences for both themselves and for humans by consuming agricultural crops. It is unclear whether savanna elephant crop consumption is merely opportunistic behaviour or related to insufficient quality of natural forage. We analysed the role of vegetation quality on elephant crop consumption. We focused on the role of micronutrients, as natural elephant diets are thought to be insufficient in elements such as sodium and phosporus, which can influence their foraging decisions.
2. For 12 months across four seasons we collected elephant feeding trail data along with tree, grass and crop samples. We investigated how the quality and availability of these items influenced elephant dietary choices across months and seasons. Subsequently, we compared levels of fibre, digestible energy, dry matter intake, and micronutrients, together with secondary compounds (tannins) across the three vegetation groups. As elephants do not make dietary choices based on one component, we also analysed the nutrient balance of food items with right-angle mixture models.
3. The levels of phosphorus, magnesium and dry matter intake corresponded to foraging preference. Compared to trees and grasses, crops contained significantly higher amounts of digestible energy content, dry matter intake, nitrogen, phosphorus, calcium and magnesium. PCA results showed that crops differed in phosphorus and magnesium levels. The right-angle mixture models indicated that except for one tree species, all food items elephants consumed were relatively deficient in phosphorus.
4. The combined results of these analyses suggest a phosphorus deficiency in elephant diet in northern Botswana. Crops, with their high absolute phosphorus levels and dry matter intake, provide an alternative source of phosphorus to reduce the deficiency. This may explain the high intensity of crop consumption in the wet season in our study area. A potential mitigation measure against elephant crop consumption might be to provide supplementary phosphorus sources.

## 1. Introduction

The consumption and destruction of crops by wildlife, often described as ‘crop raiding’, can impede co-existence of wildlife and people (Nyhus, 2016). Many rodent and mammal species, such as ungulates and primates, are known to consume crops (Naughton-Treves, 1998; Pérez & Pacheco, 2006; Arlet & Molleman, 2007; Anand & Radhakrishna, 2017). Across Asia and Africa, elephants (respectively *Elephas* spp. and *Loxodonta* spp.) are also well known for their crop consumption behaviours (Sitati et al., 2003; Hoare, 2012). African elephants consume between 100 - 300 kg of wet mass vegetation per day (Laws, 1970) and are generalist, mixed-feeders, being both browsers and grazers (Codron et al., 2011). Cultivated crops are commonly included in the diet of elephants that roam human inhabited areas (Sitati et al., 2003).

Crop consumption by elephants can threaten food security for people (Mackenzie & Ahabyona, 2012). The sharing of limited resources, and the associated close proximity of humans and elephants, can also result in conflicts causing deaths and injuries to both species (Sitati et al., 2003; Galanti et al., 2006; Kioko et al., 2008; Le Bel et al., 2010). Elephant crop consumption is prevalent in areas with high concentrations of both elephants and subsistence farmers, as in northern Botswana (Osborn, 2004; Pozo et al., 2017; Songhurst, 2017). Here, entire harvests can be destroyed by elephants, posing a threat to the livelihoods, food security, and nutrition of farmers (Gupta, 2013). As a response, the Government of Botswana and nongovernmental organizations, such as Ecoexist, partner with farmers on agricultural, policy, land use, and mitigation strategies, while also conducting research projects, including this study. Mitigation efforts often address the symptoms of crop consumption, by aiming to find ways to keep elephants out of fields. However, it is imperative that management strategies also focus on the causes of the behaviour and thus the reasons why elephants include crops in their diet (Barnes, 2002; Jackson et al., 2008). Therefore, we aim to increase understanding in how elephants make foraging decisions, and gain insights in the reasons behind crop consumption.

Given their large size, elephants are expected to be flexible in their dietary decisions, as according to the Jarman-Bell principle larger herbivores have higher digestive efficiency and a high tolerance to low quality forage (Bell, 1971; Jarman, 1974; Müller et al., 2013). Indeed, elephants do not show preferences for specific grass species, as they consume them relative to their availability (De Boer et al., 2000). However, elephants feed selectively on woody species available, neglecting or rejecting abundant forage species, and this selectiveness varies across seasons (De Boer et al., 2000; Kos et al., 2011; Owen-Smith & Chafota, 2012). Elephants also show low tolerance of secondary chemicals such as tannins and tend to avoid phenolic-rich leaves that smaller ruminants eat (Owen-Smith & Chafota, 2012). Plants develop chemical defences as tannins and other secondary chemicals to deter animals from consuming them (Molyneux & Ralphs, 1992; Kanallakan et al., 2005). These chemical plant defences are particularly present in areas with nutrient-deficient soils such as in northern Botswana (Owen-Smith & Chafota, 2012).

Seasonal patterns in crop consumption indicate that both crop and natural forage (i.e. browse and grass) quality and quantity could play a role in driving crop consumption patterns (Chiyo et al., 2005; Rode et al., 2006). During the dry season grass matures and decreases in nutritional quality, while the quality of browse changes due the availability of flowers, fruits and young leaves (De Boer et al., 2000; Kos et al., 2011; Pretorius et al., 2012; Shannon et al., 2013). During this time we expect elephants to transition from grazing to browsing, or crop consumption, as this is an attractive alternative to browsing (Osborn, 2004). Temporal variation in crop consumption correlates with crop availability at certain phenological stages (Sukumar, 1990; Tchamba, 1996; Chiyo et al., 2005; Sitati & Walpole, 2006). Agricultural crops offer high intake rates, retain high micronutrient value and a low fibre content at maturation, and contain few chemical or physical defences (Sukumar, 1990; Osborn, 2004). Therefore, elephant crop consumption is in line with predictions derived from the optimal foraging theory, selecting the best available food items from a set of foraging alternatives, based on the gain and costs of each choice (Krebs, 1977; Stephens & Charnov, 1982; Lambert & Rothman, 2015). In particular, the high levels of sodium and other micronutrients in crops, in combination with a high digestibility due to low fibre content and deterrent chemicals, could lead to crop-consumption behaviour (Rode et al., 2006).

It remains unclear to what extent elephants consume crops because of their high digestibility (i.e. low levels of fibres and secondary compounds) or their micronutrient content. Crops could simply be the best alternative, or a way to avoid dietary deficiencies in micronutrients to which elephants may be prone (Chiyo et al., 2005; Rode et al., 2006). Elephants show potential for micronutrient deficiencies (Weir, 1969; Sukumar, 1990; Holdø et al., 2002), as also illustrated by the occurrence of diseases associated with micronutrient deficiencies (Wang et al., 2007). Nutrients in which elephants are potentially deficient are sodium (Na), phosphorus (P), nitrogen (N), potassium (K), magnesium (Mg) and calcium (Ca) (Pretorius et al., 2012). Elephants can obtain their required nutrients through water sources, by geophagy – the consumption of soil (Klaus et al., 1998; Holdø et al., 2002), e.g. from termite mounds or salt deposits in caves (Weir, 1969; Bowell et al., 1996) or through optimal foraging decisions (Pretorius et al., 2012). In Kibale National Park, Uganda, forest elephants (*Loxodonta cyclotis*) are limited by minerals, rather than other factors such as energy and protein (Rode et al., 2006). Agricultural crop availability appears the main motivation for forest elephant crop consumption, while it is suggested that in savanna habitats seasonal fluctuations in natural forage quality, and therefore the risk of nutrient deficiency, may play a more important role (Chiyo et al., 2005).

To examine this hypothesis, we analysed year-round levels of micronutrients, tannins, and fibre measures (i.e. digestible energy and dry matter intake) of browse, grass and crop included in elephant diet. First, we analysed how they influenced elephant foraging choices in browse over the year (De Boer et al., 2000; Kos et al., 2011; Owen-Smith & Chafota, 2012). Secondly, we compared the levels of the vegetation quality measures between the crops, trees and grasses in order to examine whether crops are the optimal forage alternative. Finally, since animals do not make their dietary choices based solely on individual nutrient levels, but need to maintain the intake of multiple nutrients at the same time, we analysed elephant foraging options with Right-Angle Mixture Triangles (RMTs; Simpson & Raubenheimer, 1993). The use of RMTs has proven useful in understanding the dietary choices animals make (Chambers et al., 1995; Hewson-Hughes et al., 2013; Raubenheimer et al., 2015; Cabana et al., 2017). This method from nutritional geometry considers dietary choices to be based on the levels of multiple elements (Raubenheimer & Simpson, 2003; Simpson et al., 2004; Raubenheimer et al., 2014). We combined these methods to understand to what extent crop consumption is influenced by nutrient deficiencies or by opportunistic foraging behaviour.

## 2. Methods

### 2.1 Study site

We studied the role of crop consumption in the diet of elephants in the eastern panhandle of the Okavango Delta (Figure 1), an area of approximately 8,000 km^2^ in northern Botswana (Songhurst et al., 2015a). The soil in the area mainly consists of nutrient-poor Kalahari sands (Dougill & Thomas, 2004).

**Figure 1.**
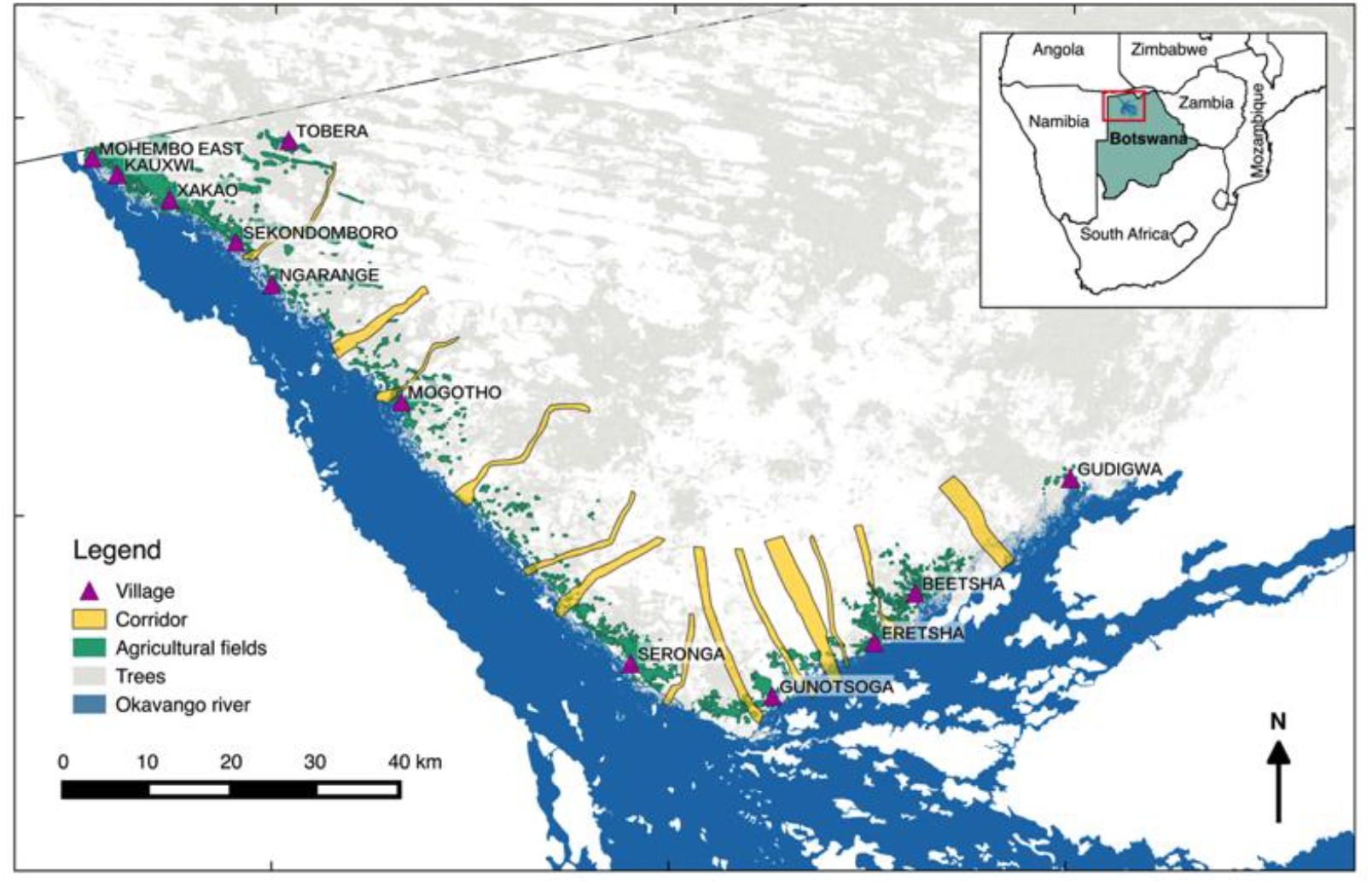
Study site in the eastern panhandle of the Okavango delta, Botswana including habitat features used in this study. The green areas represent agricultural fields, with the purple triangles representing the location of the villages, with highly used elephant corridors in yellow markings running through them towards the Okavango River in blue. White and grey areas represent savanna and tree groups, respectively.

Annually, there is one main wet season, which has on average 503 mm of rain divided over the early wet season from November until January, and the late wet season from February until April (Statistics Botswana, 2016). The crop season starts with the germination of crops in January and crop maturation continues until harvest in April-May (Songhurst, 2012). From May until July the weather becomes dry, with the late dry season from August until October. Mean maximum temperatures vary over these seasons from 26ºC in July to 36ºC in October (2000-2015, Statistics Botswana, 2016). Within this area live approximately 18,000 elephants (Songhurst et al., 2015a).

The area consists of floodplains, dry bush and agricultural fields, with seven distinguishable tree vegetation categories (Ben-Shahar, 1993; Songhurst, 2012). Around the river the floodplain vegetation type occurs, with floodplain grassland and riverine woodland with large fruit-bearing trees and small shrubs. Parts of the dry bush are dominated by mopane trees (*Colophospermum mopane*), heavily browsed by elephants, sometimes combined with other species into mixed mopane woodland. The dry bush also consists of acacia woodland with thorny browse species, areas with mixed silver terminalia (*Terminalia sericea*) vegetation, and the false mopane (*Guibourtia coleosperma*), Zambezi teak (*Baikiaea plurijuga*) and wild syringa (*Burkea Africana*) woodland. During the wet season grasses occur throughout the dry-bush area. In this area 106 pathways were identified that elephants use to walk from the uninhabited area through human dominated areas towards the river (Songhurst et al., 2015). In the same area along the river and Delta there are 13 villages with around 16,000 inhabitants (CSO, 2011). Around these villages there are (subsistence) agricultural fields including cereals like millet, sorghum and maize (*Pennisetum glaucum/ Eleusine coracana, Sorghum bicolour*, *Zea mays*, Songhurst, 2011; Heath & Heath 2009).

### 2.2 Data collection

Data were collected with permission of the Republic of Botswana Ministry of Environment, Wildlife and Tourism, research permit EWT 8/36/4 XXXI (49). First, we constructed a vegetation focal list including the mean browse and grass species included in the local elephant diet. Secondly, we followed fresh elephant feeding trails to record tree and shrub species available to, and consumed by, elephants and collected tree, grass and crop samples for nutritional content analyses.

#### 2.2.1 Constructing vegetation focal list

From August until September 2015 we constructed a vegetation focal list of species including elephant forage species present in the area verified with foraging evidence on one of seven elephant feeding trails (less than 24 h old) in each of the seven vegetation categories (Stokke, 1999; De Boer et al., 2000; Greyling, 2004; Chiyo et al., 2005; Rode et al., 2006; Kos et al., 2011; Owen-Smith & Chafota, 2012; Pretorius et al., 2012). Grass identification was verified at Wageningen University, and browse identifications were verified and included in a reference specimen collection at the Okavango Research Institute (ORI) herbarium in Maun.

#### 2.2.2 Acceptance and availability plots

From October 2015-September 2016 we followed 7-10 fresh (with spoor less than 24 h old) elephant feeding trails, during the first week of each month for 11 months (excluding April due to logistical reasons), between 6.00 AM and 6.00 PM. We took a stratified random selection of seven of the 106 pathways to focus our search effort for fresh spoor, spread over the entire region, and incorporating all dominant vegetation types in the area. We collected feeding trail information following acceptance and availability methods developed and adapted by Owen-Smith and Cooper (1987), Stokke (1999), and Greyling (2004). At the first tree with fresh elephant impact, we created a 5 m radius circular ‘food plot’ in which we recorded all trees > 20cm high that where available to the elephant, and those trees that were consumed by the elephant. Of each tree we recorded species, height, DBH, type of elephant impact (no damage, leaves removed, twigs and leaves removed, branch broken, debarked, main stem broken, uprooted) and percentage of the tree impacted. We repeatedly continued 50 m along the feeding trail and collected another food plot until in every feeding trail we collected six food plots. At every other food plot we created a ‘control plot’ similar to the food plot but 50 m perpendicular to the feeding trail, randomly to the left or right, in order to record available trees in close proximity to the feeding trail. We followed a total of 103 feeding trails, 74 from females in breeding herds and 27 from male elephants. We aimed to collect equal amounts of samples from female and male elephants but this was not feasible, as male elephant spoor was harder to find. We collected information on 594 food plots and 293 control plots. Each of these plots contained approximately 13 trees; as a result we measured 13,461 trees in total, of which 9,017 were in food plots and 4,444 in control plots.

#### 2.2.3 Vegetation content analyses

In the last month of each season (October, January, April, and July) we collected vegetation samples of all tree species on the focal list. In each month of the crop season (February, March, April) when crops and grasses were available, we also collected samples of all grass species with signs of elephant impact and of crop species. We included 27 tree species for year-round dietary choice analyses (108 trees in total), during we compared 27 tree species (an additional 81 trees), 15 grass species (45 grass samples) and 7 crop types (21 crop samples) collected during the crop season.

Vegetation samples were air-dried in a cabinet following collection, before being dried for a further 24 h at 70°C in the laboratory. Dried samples were ground to fit through a 1 mm mesh. We extracted condensed tannins using a butanol-HCl-iron assay run with 50% aqueous acetone and measured using a spectrophotometer (Mole & Waterman, 1987). We calculated the Dry Matter Intake (DMI) of the samples by measuring the Neutral Detergent Fibre (NDF), and the Digestible Energy (DE) from the concentration of Acid Detergent Fibre (ADF) in the samples. We measured the NDF and ADF by measuring sample weight differences after subsequently applying the ANKOM Fiber Analyzer vessel according to NDF and ADF guidelines (ANKOM Technology). Finally, we measured the concentration of phosphorus (P), calcium (Ca), magnesium (Mg), potassium (K), sodium (Na), and nitrogen (N) using a continuous flow analyser after destruction of the samples with salicylic acid, sulphuric acid-selenium and hydrogen peroxide (Novozamsky et al., 1983).

### 2.3 Data analyses

For our data analyses we used R (R Core Team, 2017). To construct our study site map, we used QGIS (Quantum GIS Development Team, 2015), the Semi-Automatic Classification plugin (Congedo, 2016) and Landsat 8 data, courtesy of the U.S. Geological Survey.

#### 2.3.1 Control plots

In order to test whether control plots and food plots consisted of similar vegetation, we modelled the proportion of plots in which a browse species was present versus the proportion in which it was absent, using a generalized linear model with a binomial error structure. Plot type (control or feeding trail) and month were fitted as explanatory variables.

#### 2.3.2 Acceptance/availability indices

We used the data from the food plots to calculate an index for the availability of each browse species and an index for consumption -or acceptance-of each species, per season and averaged over feeding trails. We calculated the availability index by dividing the frequency at which a species was present with the number of food plots, per season, and the acceptance index by dividing the frequency a species was accepted by their availability to elephants, per season. Plotting these acceptance and availability indices against each other for the four seasons (early dry, late dry, early wet, late wet) reveals the feeding trail-based foraging preferences and avoidances (Greyling, 2004; Owen-Smith & Chafota, 2012).

#### 2.3.3 Analysing browse choices

To examine how elephants’ dietary choices were influenced by vegetation characteristics, we constructed a generalized linear model with a binomial error structure with the seasonal proportion a species was accepted and those in which it was present as the response variable. To remove pseudo-replication, we averaged the acceptance ratio’s and vegetation characteristics over the food plots per feeding trail. As explanatory variables, we used the following vegetation characteristics: nutrient concentrations (N, Na, P, K, Mg, Ca), tannin levels, digested energy and dry matter intake percentages. The latter were based on respectively ADF and NDF percentages. NDF is a measure of the fibre content of a plant, and a plant’s NDF content determines how much of the plant an elephant can consume (van Soest & McQueen, 1973; van Soest, 1978). We use these NDF levels to calculate the daily Dry Matter Intake (DMI) for elephants: % *of DMI per kg body mass* = 120/% *NDF*, for each of the vegetation samples (Moore & Undersander, 2002). ADF can be used to calculate proxies for the energy content of vegetation. We used the ADF levels to calculate digested energy for elephants, following the formulas used by Pretorius et al. (2012): *digested energy* = 64.850 × *digested ADF*^−0.205^, *with digested ADF* = 6.665*e*^0.0246(*ADF diet*)^. Models were simplified using a backward selection procedure until variable removal significantly reduced the variance explained by the model.

#### 2.3.4 Comparing vegetation characteristics between vegetation types

We compared these same characteristics with one-way ANOVAs between trees, grasses and crops during the early (February), mid (March) and late (April) crop season, as crops and grasses are predominantly present in these months. If residuals were not normally distributed or we observed evidence of heteroscedasticity, we used non-parametric Kruskal-Wallis tests. Due to the high number of dimensions and complexity of relationships, we combined this with Principal Component Analyses (PCA) in order to visualise these differences.

#### 2.3.5 Right-angle mixture model

When making dietary decisions, animals do not only aim to avoid nutrient deficiencies, but also nutrient excess, resulting in a rule of compromise balancing the under and over consumption of nutrients (Raubenheimer & Simpson, 1999; Simpson et al., 2004). We can analyse this nutrient balance visually by plotting the different relative nutrient levels in a multidimensional space in which we plot both required and available food items (Simpson & Raubenheimer, 1993). This results into a Right-angle Mixture Triangle (RMT); a two-dimensional plot with three axes, each representing a vegetation quality measure (e.g. sodium). These axes show the percentages in which different components are present in a dietary composition (Raubenheimer, 2011). If each of the elephant food items is considered to be a composition of the elements of these three axes (e.g sodium, phosphorus, magnesium), it is possible to calculate their relative percentage based on their concentration in the vegetation samples. We calculated the ideal nutrient balance as a nutrient space from estimated minimum and average elephant dietary requirements (phosphorus: %P_min_=0.15, %P_av_=0.2, potassium: %K_min_=0.5, %K_av_=0.7, magnesium: %Mg_min_=0.1, %Mg_av_=0.3, Pretorius et al., 2012). RMTs demonstrate how balanced different food items are in their micronutrient composition, and how the elephant could combine food items to achieve the balanced diet and reach minimum nutrient requirements (Raubenheimer & Simpson, 1999; Simpson et al., 2004). Hence, we selected those vegetation characteristics that appeared most important in the first two analyses, and plotted these elements of the three types of vegetation against the required compositions of elephant diets. Since we are mainly interested in understanding the role and potential deficiency of micronutrients in elephant diet, we focus on these instead of macronutrients as conventional in RMT analyses (Raubenheimer & Simpson, 1999; Raubenheimer et al., 2015).

## 3. Results

### 3.1 Summary of results

The analyses of browsing preference indicated that elephant foraging choices are positively associated with magnesium, phosphorus, and dry matter intake. The subsequent analyses comparing vegetation types revealed that crops have higher concentrations of most nutrients, digestible energy and dry matter intake and a lower tannin concentration than browse. Finally, right-angle mixture triangles showed that elephant diet is unbalanced in phosphorus. We now explain how these findings emerged from the statistical analyses we conducted.

### 3.2 Dietary choices

#### 3.2.1 Control plots

Over all feeding trails and months, food plots had a 9% higher occurrence of most common vegetation species compared to control plots (Linear model, F_1,532_=19.58, p<0.0001).

#### 3.2.2 Acceptance/availability plots

The acceptance and availability plots reveal that changes in elephant selection of browse species are related to vegetation quality changes, as these changes occurred over the dry and wet seasons (Figure 2).

**Figure 2.**
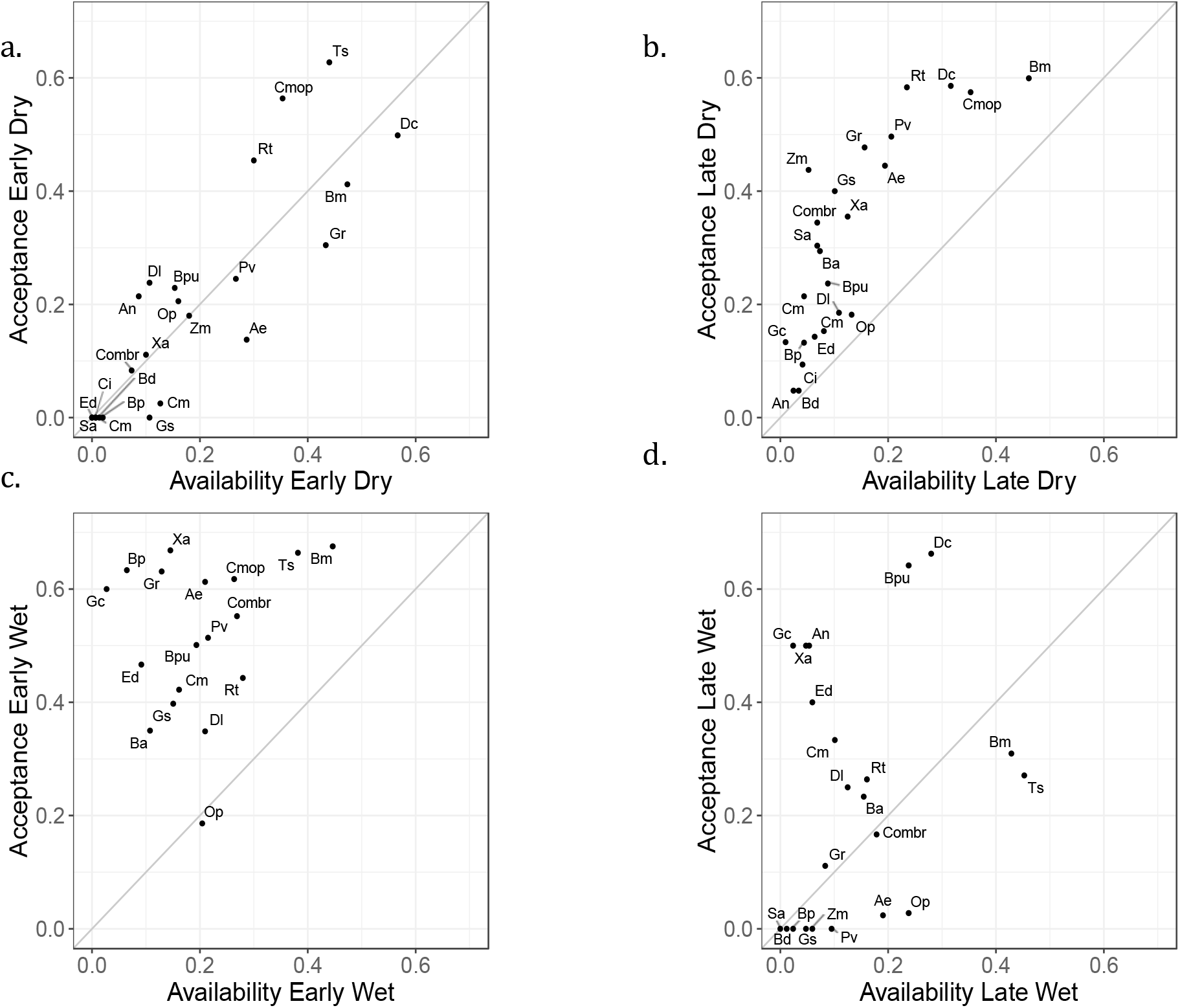
Acceptance versus availability plots divided in different seasons: a. early dry, b. late dry, c. early wet and d. late wet. Species abbreviations: Bm= Baphia mossaiensis, Ts=Terminalia sericea, Cm=Croton megalobotrys, Dl=Diospyros lycioides, Rt= Rhus tenuinervis, Op= Ochna pulchra, Ba= Burkea africana, Gs= Gymnosporia senegalensis, Gc= Guibourtia coleosperma, Cmop= Colophosperum mopane, Combr= Combretum spp., Gre= Grewia spp., Xa= Ximenia Americana/caffra, Ae= Acacia erioloba, Bpu= Bauhinia petersiana, Ed= Euclea divinorum, An= Acacia nigrescens, Cm= Combretum mossambicense, Zm= Ziziphus mucronata, Bd= Berchemia discolor, Ci= Combretum imberbe, Bp= Baikiaea plurijuga, Pv= Philenoptera violacea, Sa= Senegalia ataxancantha.

Species above the 1:1 line irrespective of season, i.e. preferred species, were *Terminalia sericea*, *Colophospermum mopane*, *Ximenia Americana/caffra*, *Guibourtia coleosperma*, *Acacia nigrescens*, *Burkea africana* and *Rhus tenuinervis*. Species for which elephants show a moderate to low preference were *Diospyros lycioides*, *Euclea divinorum*. For species such as *Dichrostachys cinerea*, *Combretum mossambicensis*, *Ziziphus mucronata*, *Baikiaea plurijuga* and *Grewia* spp. elephants show preference in some seasons, while in other seasons elephants avoided them. Elephants either avoided *Ochna pulchra*, or consumed it relative to the availability of the species. Note that in some seasons, our feeding trails did not include sufficient quantities of each species to include them into our analyses. Not only did the level of selection by elephants change, so did the general patterns of species on the plots of Figure 2. During the early dry season elephants had few preferred tree species, as most species were grouped along the line or even below it, revealing aversion. In the late dry season elephants start to show clear preference and avoidance for certain species. This preference becomes more pronounced in the early wet season, but during the late wet season elephant general tree preferences become less strong, returning to the 1:1 ratio line.

#### 3.2.3 Explaining browse choices

Elephant browse choices were influenced by season, the levels of phosphorus, magnesium and potassium, and the dry matter intake. The number of plots in which a species was eaten versus the number of plots in which it was not eaten per feeding trail, differed significantly between seasons (GLM binomial logistic regression: 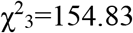, P<0.0001, parameter estimates: Early Dry µ=-2.61, SE=0.42; Late Dry: µ=0.48, SE=0.18; Early Wet µ=0.50, SE=0.19; Late Wet µ=-0.50, SE=0.22). This corresponds to the acceptance/availability plots in Figure 2, where there is a clear difference in pattern across the seasons in acceptance and availability. During the early wet and late dry when elephants show strongest species preferences, these parameter estimates are positive, while during the early dry and late wet parameter estimates are negative, corresponding to elephants being more selective and avoiding certain species. Of the nutrient levels we included in the initial model, only phosphorus, potassium and magnesium remained in the final simplified model. Of these, phosphorus and magnesium were significant in explaining the variance in the data (GLM binomial logistic regression, phosphorus: 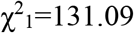, P<0.0001, parameter estimates µ=5.53, SE=1.08; magnesium: 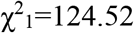, P<0.05, parameter estimate µ=0.34, SE=0.16). In particular, phosphorus was important in determining the ratio between numbers of plots where a tree is eaten compared to not eaten, with a strongly significant positive parameter estimate. On the contrary, potassium had an opposite effect with a negative parameter estimate, yet this effect was not significant (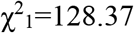, P=0.10, parameter estimates µ=-0.40, SE=0.16). Finally, dry matter intake, which is the variable we calculated based on the NDF content of the vegetation, had a positive influence on the eaten/not eaten ratio in this final model (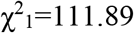, P<0.0001, parameter estimate µ=0.56, SE=0.16).

### 3.3 Comparing vegetation characteristics between vegetation types

The three vegetation types of grasses, trees, and crops differed on each of the vegetation characteristics when averaged across the crop-growing season, and crops particularly distinguished themselves from the other vegetation types by their high DMI, phosphorus and magnesium values. During the crop season, there was a steady increase in both ADF and NDF for grasses, while crops and trees remained relatively stable in their levels (ANOVA: NDF: 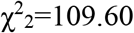, p<0.0001, ADF: 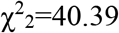, p<0.0001). Regardless of their phenological state throughout the season, crops scored highest for digestible energy content, and dry matter intake (DE: ANOVA, 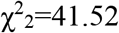, p<0.0001, DMI: Kruskal-Wallis test, F_2_=156.52, p<0.0001, Figure 3).

**Figure 3.**
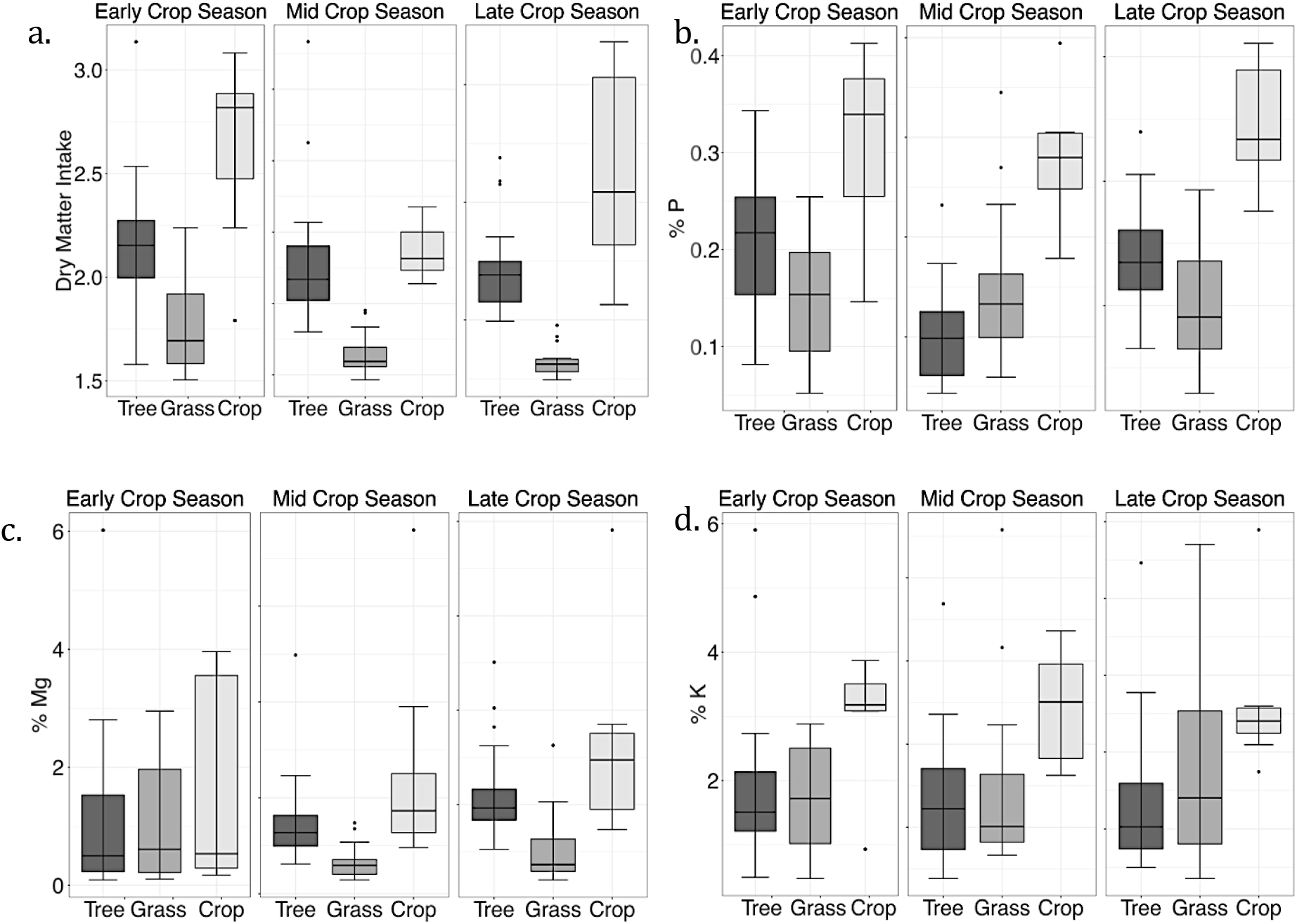
Boxplots comparing the differences in the vegetation characteristics a. Dry Matter Intake, b. Phosphorus (P), c. Magnesium (Mg) and d. Potassium (K) between trees, grasses and crops, and their changes over the crop season.

In the late crop season (April) just before crops are harvested, crops had higher concentrations of nitrogen, phosphorus, calcium and magnesium compared to trees and grasses (ANOVA: nitrogen: 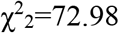, p<0.0001, phosphorus: 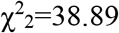, p<0.0001, Kruskal-Wallis test: potassium: 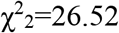, p<0.0001, calcium: 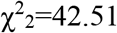, p<0.0001, magnesium: 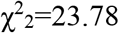, p<0.0001, sodium: 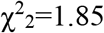, p=0.4877). During the different phenological crop stages, this difference between crops and the other vegetation types increased for calcium, magnesium and phosphorus, while it remained stable for nitrogen. The potassium levels in grasses were similar to crops, while sodium levels were highest for grasses, and maturation reduced sodium levels in crops. Tannin levels were over ten times higher for trees than for crops and grasses, and the levels remained stable across seasons (Kruskal-Wallis test: tannin: 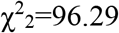, p<0.0001).

In the early crop season the first three components of the PCA explained 80% of the total variation. The first component explained over half of the variance in the data, and was loaded with each of the vegetation characteristics besides tannin. For this first component, especially digestible energy appeared to have a positive correlation, while % ADF had a similar negative influence. The second components appeared to be dominated negatively by tannin and positively by phosphorus (Figure 4). During the mid crop season, there was a small change in the percentages of variance explained in the first and second component, while the construction of the components remained the same as in the early season. During the late crop season when crops were maturing, the first component of the PCA increased again in the percentage of variance it explains, while the second reduced. The first component did not change, except tannin was no longer loaded, yet the reloading on the second component changed considerably. Nitrogen, which at first played only a role in the first component, loaded most strongly, and phosphorus remained important in the first component yet strongly reduced in its importance for the second component.

**Figure 4.**
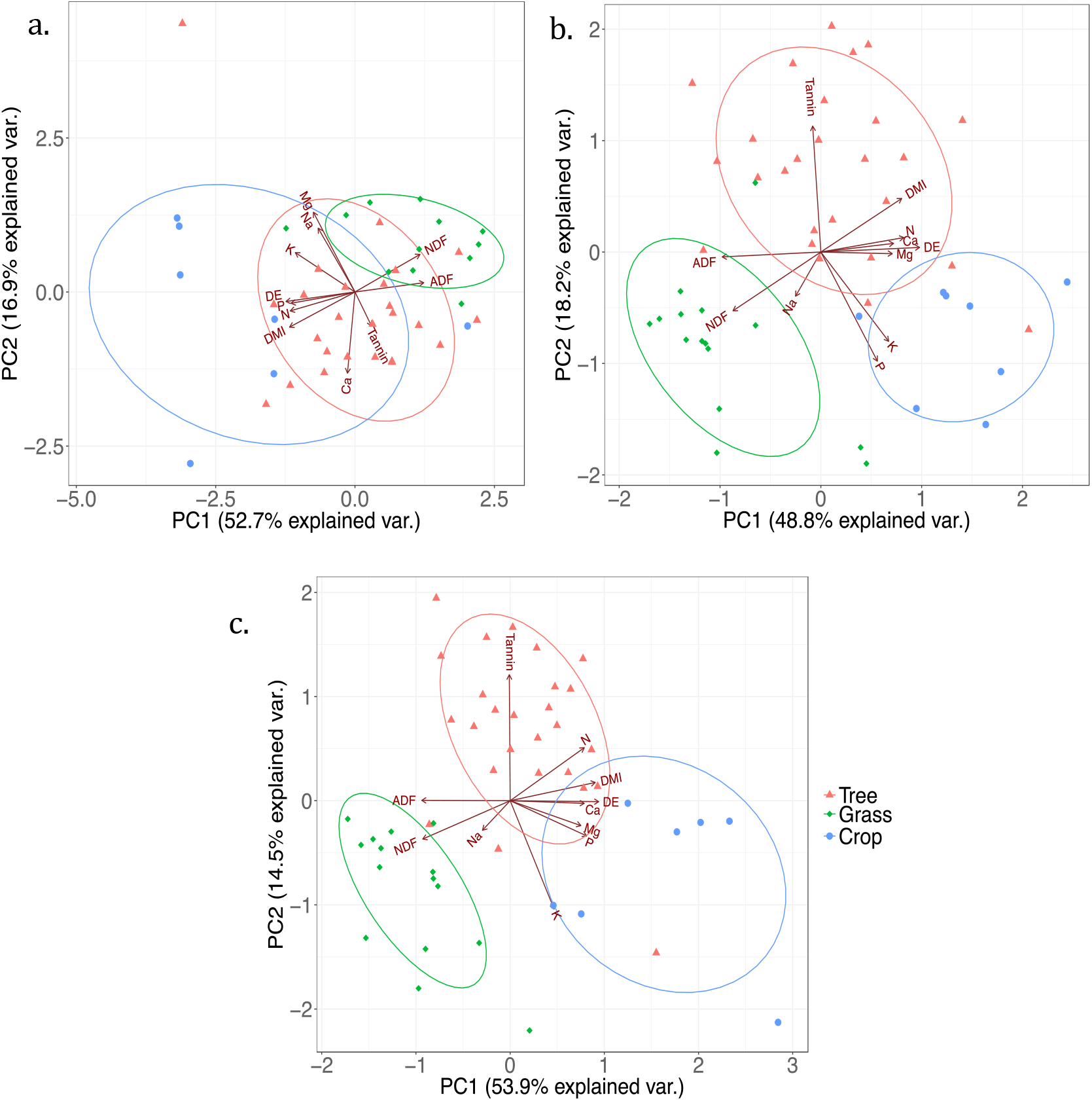
Biplots of PCAs for the a. early, b. mid and c. late crop season, revealing the clusters of groups of trees, grasses and crops.

The PCA biplots (Figure 4) showed three distinctive groups; with the grasses data grouped around the fibre measurements ADF and NDF, whereas crops were grouped among most of the nutrient variables. Sodium (Na) was more associated with grasses than with crops. Finally, trees were centred in the middle, grouped around tannin. With progression of the crop season and rainfall stimulating crop and grass growth, the differences between trees, grasses and crops increased. Crops seemed to be centred and especially distinguished from the other two vegetation types by their higher levels of phosphorus and magnesium.

### 3.4 Right-angle mixture models

Since our final model on foraging preferences included phosphorus (P), magnesium (Mg) and potassium (K), we used these three micronutrients as our RMT axes, with P on the x-axis, Mg on the y-axis and K on the tertiary axis. Because P and Mg levels were all below 60% and K levels above 40%, our axes start at these values, with the tertiary axis of 40% at the base of the triangle, and reaching 100% in the origin of the plot (Figure 5). Grey lines indicate the nutrient space between the minimal and average required nutrient balances for elephants for each of the three micronutrients. For example, food sources within the horizontal grey lines constitute the required concentration of Mg, while those above the lines have a relative surplus of Mg and those below a relative deficiency in Mg. The parallelogram created by these six grey lines represents the nutrient space in which food items are optimally balanced in these three micronutrients. If a food source lies within this nutrient space, elephants can reach a diet balanced in these three micronutrients by only consuming that food item. Nevertheless, it is also possible to reach this dietary balance by combining food sources, and thus consuming food items with matching surpluses and deficiencies in other to reach a balance on average (Raubenheimer et al., 2015). Our RMT plots indicate that the ratio between P:Mg:K varies over the seasons, with in the early and mid crop season excessive ratios of Mg and K, and in each of the seasons a relative imbalance in phosphorus (Figure 5). Only one tree species *Ochna pulchra* reached the required P level in the late crop season with 19%. In the late crop seasons when crops are mature, the P:Mg:K ratios of trees, crops and grasses are clustered. We also display these plots including the dry matter intake levels based on the NDF (Figure 6a) and relative condensed tannin (Figure 6b).

**Figure 5.**
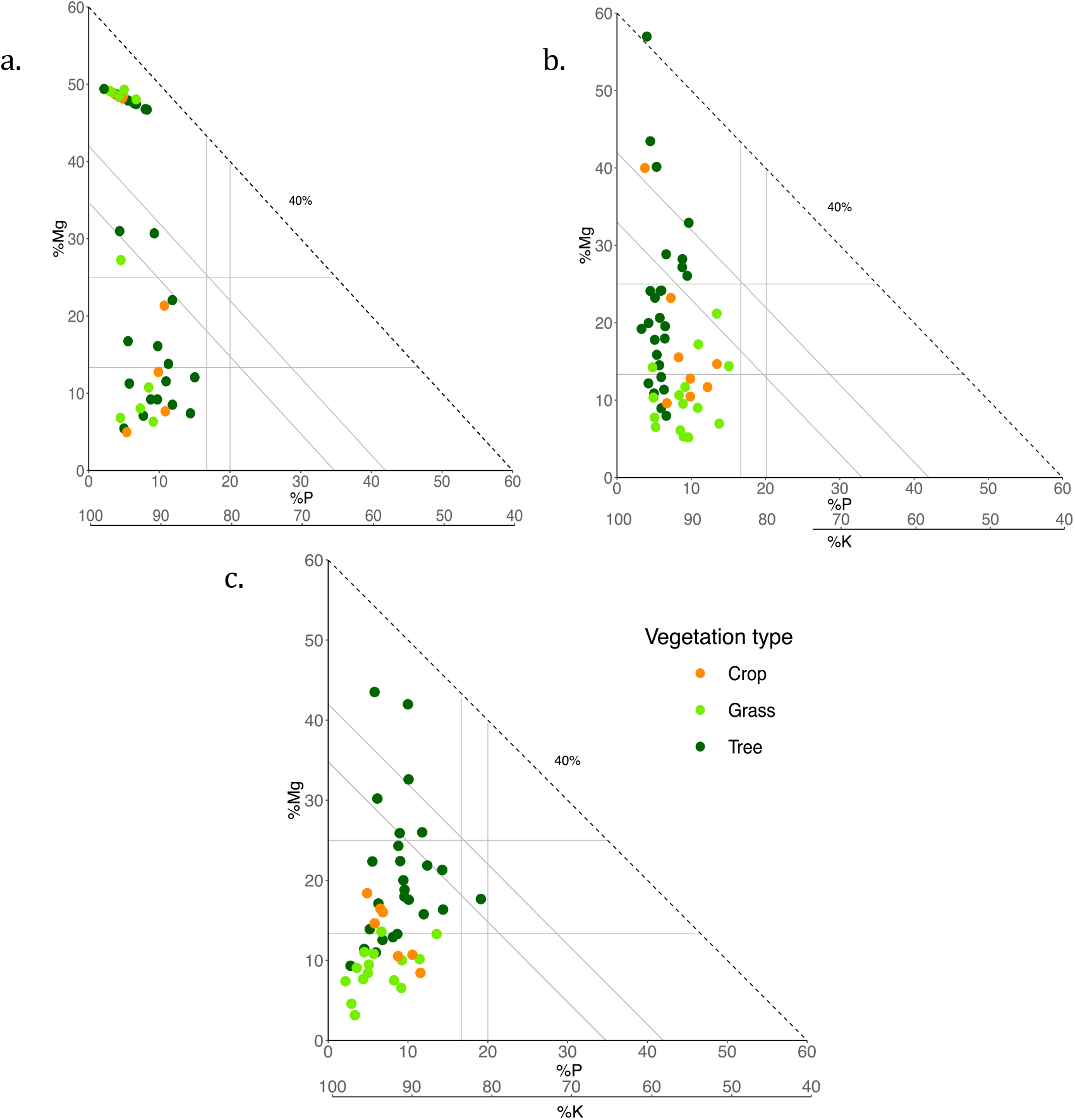
Right-angle Mixture Triangles, plotted with phosphorus on the X-axis (P), magnesium on the Y-axis (Mg) and potassium on the diagonal Z-axis (K). Plot a, b and c show the P:Mg:K ratio for respectively the early, mid and late crop season, for crops, grass and trees. The axes are scaled from 0-60% and 100-40%, none of the points contained a percentage of Mg or P higher than 60%, and only one a percentage of K lower than 40%.

**Figure 6.**
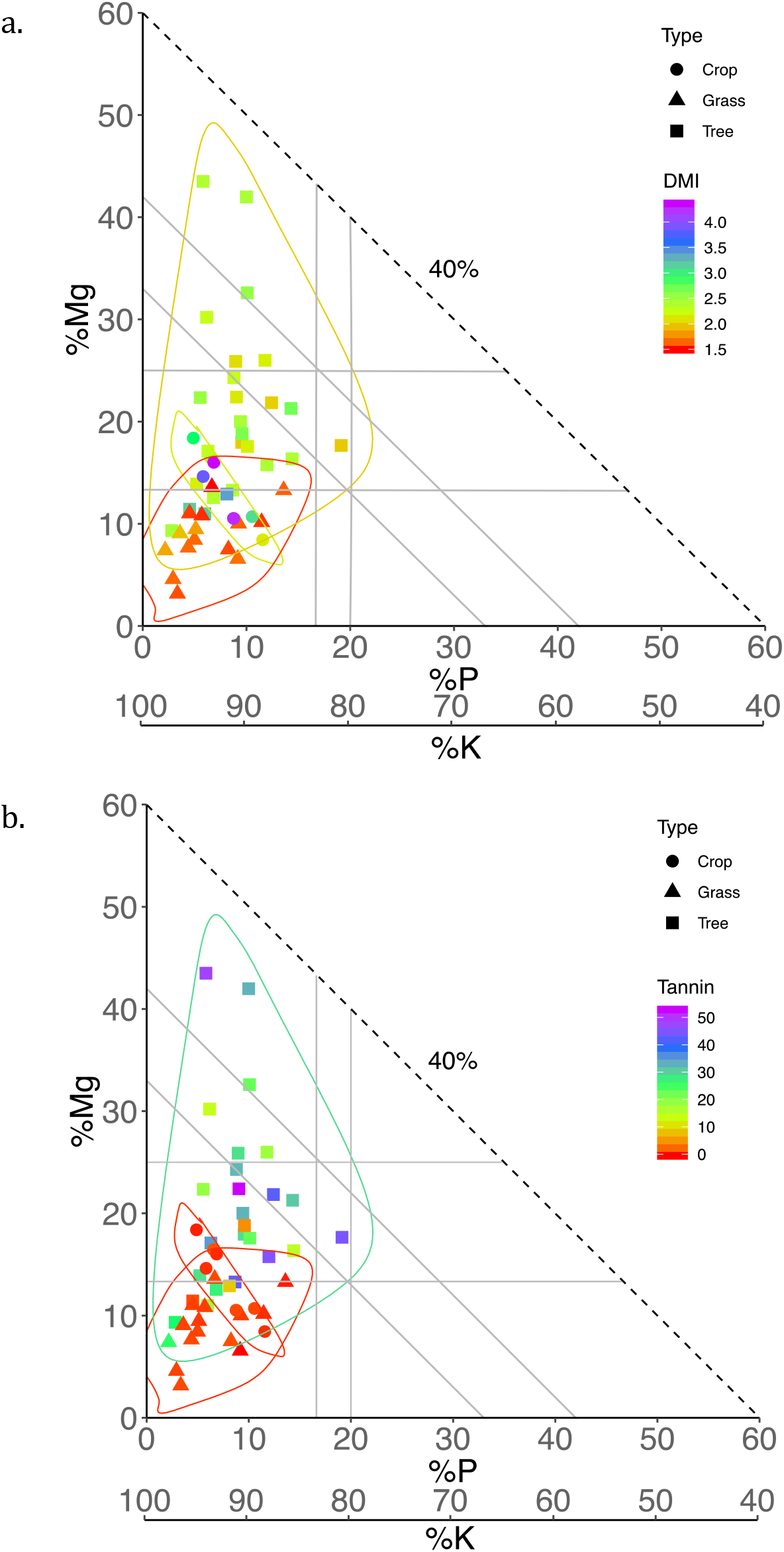
Plots shows the same data as plot c in Figure 5, however the colour of the food items indicates the dry matter intake levels (a) or condensed tannin (b). The axes are scaled from 0-60% and 100-40%, none of the points contained a percentage of Mg or P higher than 60%, and only one a percentage of K lower than 40%.

## 4. Discussion

Our results suggest that in our study site micronutrient concentrations are an important determinant in elephant crop consumption, and that crop consumption could be a strategy to avoid or minimize dietary deficiencies.

The acceptance/availability plots indicate that foraging preferences vary over the season. Our analyses of foraging preference indicate that elephants select browse species based on the dry matter intake value and concentrations of phosphorus and magnesium, and potentially potassium. Phosphorus and magnesium had a positive effect on browse preference. Dry matter intake appeared to also have a positive influence on dietary preferences towards tree species. This appears contrary to previous research that showed fibre measures were unrelated to elephant food intake (Meyer et al., 2010).

Our comparison between crops, grasses, and trees on nutrient and fibre measurements showed that grasses were highest in ADF and NDF fibre contents, and that these levels increased towards maturation when the fresh green grass started to dry. By contrast, the digestible energy and dry matter intake were highest in crops; thus consuming crops increases energy levels faster than consuming grass or trees. This concurs with previous research that showed that digestible energy is an important factor in elephant dietary optimisation (Pretorius et al., 2012). Tannin levels were significantly higher in trees than in crops and grasses, making them less desirable for digestion (Owen-Smith & Chafota, 2012). However, in our analysis tannin levels did not influence elephant dietary browse preferences, suggesting that unless there is a threshold relationship above which tannins do not play a role, tannin levels are not an important driver in forage choice by elephants. This could be related to the large salivary glands which may buffer against the effect of tannins (Schmitt, 2017). Even if there is a threshold relationship, tannin cannot explain elephants consuming crops over grasses, as there was no significant difference between the tannin levels of crops and grasses.

During crop maturation, nutrient concentrations in crops became significantly higher than those in browse and grass, except for sodium, which was more available in grasses than crops. Therefore, we did not find support for sodium deficiency in elephant diet in our study area or evidence that crop consumption is a response to sodium cravings, in contrast to comparable studies in other areas (Sukumar, 1990; Holdø et al., 2002; Rode et al., 2006).

The clustering of trees, grasses and crops in the PCA concentrated into separate groups towards the end of the crop season. Nutrients played an important role in explaining the variation within the data, with crops clustered around a correlated group of dry matter intake, digestible energy, phosphorus, magnesium, calcium and potassium. Browse species were mainly concentrated around tannin, nitrogen, and grass around the fibre measures NDF and ADF and sodium levels.

Finally, the RMT graphs displayed how the ratios between the three nutrients were distributed over trees, grasses and crops. Grasses appeared to result in the highest and trees in the lowest relative potassium percentages, with crops in an intermediate position. Regarding magnesium, crops contained intermediate percentages compared to trees and grasses. At the same time, most trees achieved the required ratio in magnesium, while most grasses had lower values. While there were multiple food sources that fell within the nutrient space indicating balanced magnesium and potassium values, neither natural forage nor crops reached a nutrient balance for elephants regarding phosphorus, revealing a potential deficiency in phosphorus in elephant diet. An increase in the ratio between calcium and phosphorus could furthermore accentuate a deficiency in available phosphorus (McNaughton, 1990). The intermediate position of crops could also contribute to crops’ attractiveness to elephants. By selecting crops, elephants could balance out the excess of potassium and possibly calcium and deficiencies in other nutrients, which in the RMT framework is considered a ‘rule of compromise’ (Raubenheimer & Simpson, 1999). Moreover, the RMT plot including the dry matter values (Figure 6.a), clearly illustrates the significantly lower dry matter values of crops, meaning that elephants can consume significantly more crops than trees and grasses, thus allowing a higher possibility of consuming sufficient amounts of phosphorus. The RMT plot including tannin levels (Figure 6.b) visualises the higher tannin levels of trees, however we know from the vegetation content comparisons that there was no significant difference between tannin levels of grasses and crops.

## 5. Conclusion & management implications

Together, our results provide insights into the importance of micronutrients in crop consumption behaviour, and the potential effectiveness of mitigation measures such as artificial salt licks (Zhang & Wang, 2003). Our study suggests that consuming crops could be more than just a better alternative to browse and grass, and could even be a necessity to cope with micronutrient deficiencies in natural forage. Crops are a better option to browse and grass due to their higher dry matter intake, digestible energy and micronutrient values. However, the importance of phosphorus levels in increasing browsing preference, the extreme levels of phosphorus in crops, the importance of phosphorus in clustering the vegetation types and furthermore the potential phosphorus deficiency indicated by the RMT models, suggest that crop consuming behaviour in elephants could be explained by a phosphorus deficiency when only feeding on grasses and trees. Phosphorus has more known functions than any of the major minerals (McDonald et al., 2011) and plays an important role in the development of cells and tissues (Ihwagi et al., 2011), energy metabolism and is in close association with calcium in bone (McDonald et al., 2011). Deficiencies in phosphorous are widespread, since most soils worldwide are deficient in this element, especially in (sub-) tropical regions (McDonald et al., 2011, McDowell 2003, O’Halloran et al., 2010). Deficiencies in phosphorus can have a direct impact on fertility and reproduction (McDonald et al., 2011). Elephants can crave phosphorus, suggested to be the main reason behind tree debarking, due to the high concentrations of phosphorus in bark (Ihwagi et al., 2011). Elevated levels of phosphorus can also be found in soil licks (Klaus et al., 1998) and in vegetation on termite mounds (Grant & Scholes, 2006). Further research including absolute dietary input is needed to confirm the role of phosphorus deficiency in stimulating elephant crop consumption, taking into account not only the quality but also the quantity of forage elements. Our study also reveals the importance of including information on feeding ecology into addressing crop consuming behaviour, as these influences can be site specific. Incorporating knowledge on crop consumption drivers into mitigation measures can increase efficiency and effectiveness of them. For example, providing supplementary licks rich in phosphorus away from crop fields warrants further investigation as a crop consumption mitigation technique.

## 6. Acknowledgement

We would like to thank the Ecoexist Project, The Howard G. Buffet Foundation, and the Government of Botswana for facilitating data collection. We especially want to thank Olorato Ratama, Mpotshang Fabian France, and Rodgers Keemekae for their contributions to the data collection. We are also thankful to the NERC Oxford DTP Environmental Research, Pembroke College Oxford, and Stichting dr. Hendrik Muller’s Vaderlandsch Fonds for their financial support. We would like to thank the Okavango Research Institute, in particular Frances Murray-Hudson, Chanana Kupe, Lindah Maekopo and Joseph Madome, and Wageningen University, in particular Jan van Walsem, Herbert Prins and Elmar Veenendaal for their facilitation roles in data collection and labwork. Finally, we would like to thank members of the Department of Zoology for their helpful advice, especially Sonya Clegg, Lucy Taylor, Harriet Downey, Chris Terry, James Foley and all members of research group E2D, especially Shelly Lachish, Rosemarie Kentie, Leejiah Dorward, Erik Sandvig, and Emily Simmonds.

## 7. Data accessibility

Data will be available from the Figshare data repository.

## 8. Author contributions

SMV conceived the idea for the study and SMV, WFB and MDH designed methodology; SMV, TC, ACS, GM and ALS adapted methods to study site; SMV and MM lead data collection; SMV and TC analysed the data; SMV and TC led the writing of the chapter. All authors contributed critically to the drafts and gave final approval for publication.

