## Supplementary Information for "Do African savanna elephants (*Loxodonta africana*) eat crops because they crave micronutrients?"

**Appendix I – Study site vegetation types**

*Table 1. Study area divided into main vegetation categories defined by tree, shrub and grass species occurring in the categories.*

| Area | Vegetation category | Trees and shrubs | Grasses and plants |
|---|---|---|---|
| Floodplain | Floodplain grassland and Riverine woodland | Jackal berry (Diospyros mespiliformis), water berry (Syzyginum spp.), sausage tree (Kigelia africana), leadwood (Combretum imberbe), large fever-berry (Croton megalobotrys), marula (Sclerocarya birrea), large-fruited bushwillow (Combretum zeyheri), red star apple (Diospyros lycioides), magic guarri (Euclea divinorum), brown ivory (Acacia erubescens), knobbly combretum (Combretum mossambicense), white bauhinia (Bauhinia petersiana), kalahari currant (Commiphera rhus), rough leaved raisin (Grewia flavescens), shepard’s tree (Boscia albitrunca), russet bushwillow (Combretum hereroense), sickle-leaved | Swamp savanna grass (Miscanthus Junceus), mat sedge (Schoenoplectus corymbosus), African bristlegrass (Setaria sphacelata), drop seed (Sporobolus fimbriatus), couch grass (Cynodon dactylon), phuka (Urochloa brachyuran/trichopus), false signal grass (Brachiaria deflexa), torpedograss (Panicum repens) |

| | | | |
|---|---|---|---|
| | | albizia (Albizia harveyi), confetti tree (Gynmosporia senegalensis), sourplum spp. (Ximenia americana, caffra), raintree (Philenoptera violacea), buffalo thorn (Ziziphus mucronata), peeling bark (Ochna pulchra) | |
| Dry bush | Silver terminalia sandveld | Silver terminalia (Terminalia sericea), sand camwood (Baphia massaiensis), mopane, acacia species, rain tree (Philenoptera violacea), white bauhinia (Bauhinia petersiana), kalahari current (Commiphera rhus), rough leaved raisin (Grewia flavescens), shepard's tree (Boscia albitrunca), marula (Sclerocarya birrea), large-fruited bushwillow (Combretum zeyheri), russet bushwillow (Combretum hereroense), confetti tree (Gynmosporia senegalensis), sickle bush (Dichrostachys cinerea), raintree (Philenoptera violacea), Camel thorn (Acacia erioloba), knobthorn | *For all dry bush categories:* Couch grass (Cynodon dactylon), phuka (Urochloa brachyuran/trichopus), false signal grass (Brachiaria deflexa), torpedograss (Panicum repens), silky bushman grass (Stipagrostis uniplumus), lovegrasses (Eragrostis porosa, Eragrostis rotifer, Eragrostis lehmaniana) |

| | | | |
|---|---|---|---|
| | | (Acacia nigrescens), peeling bark (Ochna pulchra) | |
| | Mopane woodland | Mopane (Colophospermum mopane), sand camwood (Baphia massaiensis) | |
| | Mixed mopane woodland | Mopane (Colophospermum mopane)* | |
| | Acacia woodland | Camel thorn (Acacia erioloba), knobthorn (Acacia nigrescens), flame thorn (Senegalia ataxacantha), buffalo thorn (Ziziphus mucronata)* | |
| | False mopane, Zambezi teak and wild syringa woodland | False mopane (Guibourtia coleosperma), Zambezi teak (Baikiaea plurijuga), wild syringa (Burkea africana), peeling bark (Ochna pulchra), sand camwood (Baphia massaiensis) | |
| Agricultural fields | Crops | | Millet (Pennisetum glaucum/ Eleusine coracana), sorghum (Sorghum vulgare) and maize (Zea mays), beans (Vigna |

aconitifolia/ Phaseolus vulgaris), groundnuts (Arachis hypogaea), watermelon (Citrullus lanatus), pumpkin (Cucurbita spp.)

* This category additionally includes the species of category ‘Silver terminalia sandveld’ in limited amounts.

### Appendix II – Functions and deficiencies of micronutrients in mammals

Nutrients in which elephants are potentially deficient are sodium (Na), phosphorus (P), nitrogen (N), potassium (K), magnesium (Mg) and calcium (Ca) (Pretorius *et al.*, 2012). In a worldwide study of nutrient deficiencies in grazers, elements that appeared to be limiting in southern Africa were Mg, P, Na, cupper (Cu), iodine (I), manganese (Mn) and selenium (Se) (Mcdowell *et al.*, 1977). However, since Mn deficiencies mainly occur in poultry this element is to our knowledge not studied in relation to herbivores (McDowell, 2003). A study of the Serengeti National Park showed that in savanna grasslands, herbivores are particularly prone to deficiencies in Mg, Na and P (McNaughton, 1988).

#### 2.1 Sodium

Sodium (Na) deficiencies are common in many parts of the world, especially in tropical areas in Africa. This deficiency causes lower osmotic pressure and dehydration of the body, resulting in poor growth, and a reduction in the utilization of protein and energy that is digested (McDonald et al. 2011). The occurrence of this deficiency is likely in the case of rapidly growing (young) animals that feed on forage low in Na, which is the case for most tropical forage. Other factors contributing to deficiency are the loss of sodium chloride (NaCl) due to sweating, lactating, and high levels of potassium (K), like in fertilized pastures, since K excess worsens Na deficiency (McDowell, 2003). It is unclear what exactly are the sodium requirements of elephants, yet, there is sufficient evidence that salt craving or sodium carving occurs in elephants and influences their behaviour (Holdø, Dudley and McDowell, 2002; Rode *et al.*, 2006). There is evidence for a naturally occurring deficiency in sodium levels in the diet of grazers, especially when lactation requires elevated sodium levels (Mcdowell *et al.*, 1977; Jachmann and Bell, 1985). Consequently a well-known example of nutrient deficiency in elephants is sodium drive or craving, which could be leading elephants to consume crops to fulfil their sodium requirements (Sukumar, 1990; Rode *et al.*, 2006). In general, crops have relatively high sodium concentrations compared to natural forage, which in light of expected deficiencies makes crop consumption highly attractive (Sukumar, 1990). Other known sodium sources are surface water bodies, and as elephants depend more on these water sources in the dry season, the demand to receive sodium through other sources - such as foraging- is less in this season (Weir, 1969; Pretorius *et al.*, 2012).

Weir (1972) discovered that there is a close correlation between the level of sodium concentration of a particular water source, and the number of elephants that make use of this source. At the same time, other sodium sources, such as ‘salt licks’ were ignored in these areas with sodium rich water. Moreover, the use of salt licks was not due to other (Weir, 1972). Chamaillé-Jammes however point out that this study took place in a period of low elephant population density. Therefore, they re-analysed the relationship between elephant number and sodium concentrations in waterholes over the period of Weir’s study and added new data periods until 2005. This study showed that indeed, the relationship was highly significant during Weir’s study period in the early 1960’s, yet this was not true for the subsequent periods, thus elephants did not favour the sodium rich water sources over others. Unfortunately it remains unclear what could motivate this change in water source selection (Chamaillé-Jammes, Fritz and Holdo, 2007).

Holdø et al. (2002) also re-examined the hypotheses of Weir (1972) that sodium drive in elephants determined their distributions, in addition to that they analysed the Na content of natural forage. Their conclusion is that during the dry season elephants in the Kalahari supplement their Na intake with ‘mineral licks’, as concentrations in vegetation are low. This means that the Na licks appear to affect their movement and habitat use. Even though salt licks also contain above average levels of Ca and Mg, it is unlikely that salt lick use could be attributed to that (Holdø, Dudley and McDowell, 2002). This co-occurring is associated with the positive connection between the concentrations of sodium and magnesium, and in turn between magnesium and calcium (Jachmann and Bell, 1985). The use of the licks increases the amount of Ca and Mg that is secreted in faeces, which makes it unattractive to elephants deficient in these minerals. Opposite, the faeces of elephants that made use of the licks appeared to have low Na concentrations, which suggests that if these elephants are indeed deficient in Na, their gut is capable of electrolyte absorption to reduce Na loss (Holdø, Dudley and McDowell, 2002).

#### 2.2 Potassium

Besides its occurrence in many studies analysing elephant nutrition, deficiencies in K levels tend to be rare in grazers since most plants have high K contents (McDonald *et al.*, 2011). Still this element often returns in studies of elephant nutrition, probably since it is one of the essential macro elements (Weir, 1972; Jachmann and Bell, 1985; Rode *et al.*, 2006; Ihwagi *et al.*, 2011; Pretorius *et al.*, 2012). Together with sodium, chlorine and bicarbonate ions it plays important roles in osmotic regulations of the body fluids, nerve and muscle system and metabolism (McDonald *et al.*, 2011). Even though this element occurs in bark and salt licks that are used by elephants, it is probably not the main motivator to consume them (Weir, 1969; Holdø, Dudley and McDowell, 2002; Ihwagi *et al.*, 2011).

#### 2.3 Magnesium

Deficiencies in magnesium are uncommon in animals and humans (McDowell, 2003). Nevertheless, the by elephants often utilized salt licks contain elevated levels of magnesium concentration (Weir, 1969; Klaus, Klaus-Hügi and Schmid, 1998; Holdø, Dudley and McDowell, 2002). It is also often included in studies, without justification (Jachmann and Bell, 1985; Sukumar, 1990; Wang *et al.*, 2007; Ihwagi *et al.*, 2011; Pretorius *et al.*, 2012). This has probably to do with the close association is has with calcium and phosphorus, and its essential importance in efficient metabolism of carbohydrates and lipids. Furthermore, magnesium content shows a high variability between different forage sources, so deficiencies do occur occasionally (McDonald *et al.*, 2011).

**2.4 Calcium**

Research in Malawi has shown that deficiencies in calcium (Ca) are common among grazers, especially in the dry season (McDowell, 2003). This occurrence of deficiencies is related to the low content of calcium in natural forage (Wang *et al.*, 2007). Calcium is the most abundant mineral element of bodies, and is important for the skeleton, teeth, living cells, tissue fluids, and the functioning of enzymes, nerves and muscles (McDonald *et al.*, 2011). Deficiencies could cause problems to elephants with their muscles, bones, eyes, and paralysis of their trunk and throat (Wang *et al.*, 2007). In contrast to wild grasses, cultivated crops often have high levels of calcium, and crop consumption in order to raise their calcium levels could therefore be an optimal foraging strategy for elephants (Sukumar, 1990; Von Gerhardt *et al.*, 2014). Besides, bark of trees is also high in calcium, yet there is disagreement on the importance of the presence of calcium on stimulating tree debarking (Barnes, 1982; Ihwagi *et al.*, 2011). Calcium also often occurs in the sodium rich soil and water consumed by elephants, yet it seems unlikely that calcium plays an important role in the existence of these behaviours (Weir, 1969, 1972). Finally, calcium is one of the nutrients that are present in the salt licks (Weir, 1969; Holdø, Dudley and McDowell, 2002).

**2.5 Phosphorus**

Deficiencies in phosphorus (P) are widespread, since most soils worldwide are deficient in this element, especially in (sub-) tropical regions (McDonald et al., 2011, McDowell 2003, O'Halloran et al., 2010). Phosphorus has more known functions than any of the major minerals (McDonald *et al.*, 2011). Phosphorus plays an important part in the development of cells and tissues (Ihwagi *et al.*, 2011), energy metabolism and is in close association with calcium in bone, while a deficiency has direct impacts on fertility and reproduction (McDonald *et al.*, 2011). Debarking of trees by elephants could be motivated by the relatively high concentrations of phosphorus in bark (Ihwagi *et al.*, 2011). Elevated levels of phosphorus can also be found in soil licks (Klaus, Klaus-Hügi and Schmid, 1998) and in vegetation on termite mounds (Grant and Scholes, 2006).

#### 2.6 Nitrogen

Nitrogen can be used to measure crude protein of vegetation, since protein consists of nitrogen, together with other organic compounds such as carbon, hydrogen and oxygen (McDonald *et al.*, 2011). Nitrogen is considered to be among the most limiting of all nutrients for in the vegetation and for herbivores in Africa (O'Halloran *et al.*, 2010; Codron *et al.*, 2011). In modelling forage selection by elephants, Pretorius et al. (2012) observed that during the wet season elephants tend to maximize their nitrogen intake (Pretorius *et al.*, 2012).

#### 2.7 Iodine

Another widespread deficiency is in the element iodine (I), which occurs especially in areas where the soil has been depleted, and rain and wind are inadequate to provide enough I from its oceanic source. Of the micronutrients, iodine is particularly important for metabolism and overall general health (McDowell, 2003). Although iodine not often is included in elephant foraging studies, Milewski (2000) argues that elephants are prone to iodine deficiency, since they will require high amounts of iodine, and their food sources are deficient in the element. This iodine craving could drive them to artificial bore water (Milewski, 2000).

### 2.8 Other micronutrients

Besides these essential major elements, there are also essential micro or trace elements: iron (Fe), iodine (I), manganese (Mn), zinc (Zn) and cobalt (Co). These minerals can be very important to the metabolism of the body, but need to be present in smaller quantities than major elements (McDonald et al. 2011). To put this in perspective; on average the body nutrients are made up of 46% Ca, 29% of P, 25% of K, S, Na, Cl and Mg, while the trace elements together contribute to less than 0.3% of the body nutrients (McDowell, 2003). Minerals are held in the central reserve in the body, usually the blood plasma or bones in the case of Ca, and interchange the minerals by secretion into other compartments (McDonald et al. 2011, McDowell, 2003).

**Appendix III – Plant species elephants included in elephant diet**

| Code | Common name | Latin name | Sestswana name |
|---|---|---|---|
| SCW | Sandcamwood | *Baphia massaiensis* | / |
| SB | Sickle bush | *Dichrostachys cinerea* | Moselesele |
| RT | Rain tree | *Philenoptera violacea* | Mopororo |
| CaT | Camel thorn | *Acacia erioloba* | Mogotho |
| Mop | Mopane | *Colophospermum mopane* | Mophane |
| SP | Sour plum spp. | *Ximenia americana, caffra* | Moretologana, Morokolo |
| ST | Silver terminalia | *Terminalia sericea* | Mogonono |
| JB | Jackalberry | *Diospyros mespiliformis* | Mokutshume |
| RSA | Red star apple | *Diospyros lycioides* | |
| ConfT | Confetti tree | *Gynmosporia senegalensis* | Mothone |
| LW | Leadwood | *Combretum imberbe* | Motswere |
| SLA | Sickle leaved albizia | *Albizia harveyi* | / |
| RLR | Rough leaved raisin | *Grewia flavescens* | Mokgompatha |
| WB | White bauhinia | *Bauhinia petersiana (urbaniana)* | Motshantsha |
| KT | Knobthorn | *Acacia nigrescens* | Mokaba |

| | | | |
|---|---|---|---|
| LFBerry | Large fever berry | *Croton megalobotrys* | Motsebi |
| Mag | Magic guarri | *Euclea divinorum* | Mothakola |
| ZT | Zambezi teak | *Baikiaea plurijuga* | |
| WS | Wild Syringa | *Burkea africana* | Mosheshe |
| FM | False Mopane | *Guibourtia coleosperma* | |
| Mar | Marula | *Sclerocarya birrea* | Marula |
| LFBush | Large fruited bushwillow | *Combretum zeyheri* | / |
| OP | Peeling bark | *Ochna pulchra* | Monyelenyele |
| KC | Kalahari currant | *Rhus tenuinervis* | Morupaphiri |
| BuffT | Buffalo thorn | *Ziziphus mucronata* | Mokgalo |
| BlueT | Blue thorn | *Acacia erubescens* | Moloto |
| ShepT | Shepard tree | *Boscia albitrunca* | Motopi |
| Russ BW | Russet bushwillow | *Combretum hereroense* | Mokabi |
| BrIv | Brown ivory | *Berchemia discolor* | Motsintsila |
| Kcomb | Knobbly creeper | *Combretum mossambicensis* | Motsheketsane |

**Appendix IV – Data collection classification categorizations**

Table 1. Description of plant elephant impact types included in the study.

| Impact type code | Description | Range of damage % |
| --- | --- | --- |
| No damage | No sign of elephant impact | 0 |
| Lv | **Leaves:** Only leaf stripping | 0-10 |
| Tw,lv | **Twigs, leaves:** Only twigs (usually <5 cm circumference) and leaves removed | 10-30 |
| Br | **Branches:** Branches are broken and/or bark stripped in most cases branches are >5cm circumference | 10-50 |
| Deb | **Debarking**: The bark is stripped from the main stem | 10-50 (unless ringed and dead than 100%) |
| MS | **Main stem broken**: The main stem of the tree is broken or removed. | 50-100 (100 if tree dead) |
| R | **Root damage/uprooting**: The elephants have dug up the roots, and/or have removed or debarked them | 10-100 (100 if completely uprooted and tree dead) |

Table 2. Description of Forage Quality Index (FQI) types included in the study.

| FQI code | Description | Range of FQI% |
|---|---|---|
| No | There is nothing on the tree | 0 |
| Old | There are old leaves on the tree | 25-100, OR <10 |
| Bud | There are leaves or flower buds on the tree, and some are starting to open | 0-25, OR <10 |
| New | The buds have opened and there are new leaves on the tree | 25-100, OR <10 |
| Fruit | There are fruits growing on the tree (fresh fruits, not seeds) | 25-50, OR <10 |
| Flower | There are flowers on the tree | 25-50, OR <10 |

**Appendix V**

Table 1. GLM with binomial error structure of the proportion of plots in which a species is eaten in which it is found in each month (d.f.=97).

| **Explanatory variable** | Estimate | Standard error | z-value | p |
|---|---|---|---|---|
| **Early Dry (intercept)** | -2.60958 | 0.42082 | -6.201 | <0.0001 |
| **Early Wet** | 0.50343 | 0.19256 | 2.614 | <0.01 |
| **Late Dry** | 0.47674 | 0.18138 | 2.628 | <0.01 |
| **Late Wet** | -0.50209 | 0.21566 | -2.328 | <0.05 |
| **% P** | 5.52599 | 1.08332 | 5.101 | <0.0001 |
| **% K** | -0.39674 | 0.15871 | -2.500 | <0.05 |
| **% Mg** | 0.34204 | 0.16203 | 2.111 | <0.05 |
| **Dry Matter Intake** | 0.5604 | 0.1579 | 3.551 | <0.001 |

Table 2. Test results of comparing fibre measurements between vegetation types.

| Explanatory variable | Df | Test | F/Chi-Square | p |
|---|---|---|---|---|
| NDF | 2 | One-Way ANOVA | 109.6 | <0.0001 |
| ADF | 2 | One-Way ANOVA | 40.39 | <0.0001 |
| Digestible Energy | 2 | One-Way ANOVA | 41.52 | <0.0001 |
| Dry Matter Intake | 2 | Kruskall Wallis | 156.52 | <0.0001 |
| N | 2 | One-Way ANOVA | 72.98 | <0.0001 |
| P | 2 | One-Way ANOVA | 38.89 | <0.0001 |
| K | 2 | Kruskall-Wallis | 26.516 | <0.0001 |
| Ca | 2 | Kruskall-Wallis | 42.511 | <0.0001 |
| Mg | 2 | Kruskall-Wallis | 23.783 | <0.0001 |
| Na | 2 | Kruskall-Wallis | 1.8489 | 0.4877 |
| Tannin | 2 | Kruskal-Wallis | 96.288 | <0.0001 |

**Appendix VI – Non-significant boxplot comparisons between tree, grass and crops and changes over the crop season.**

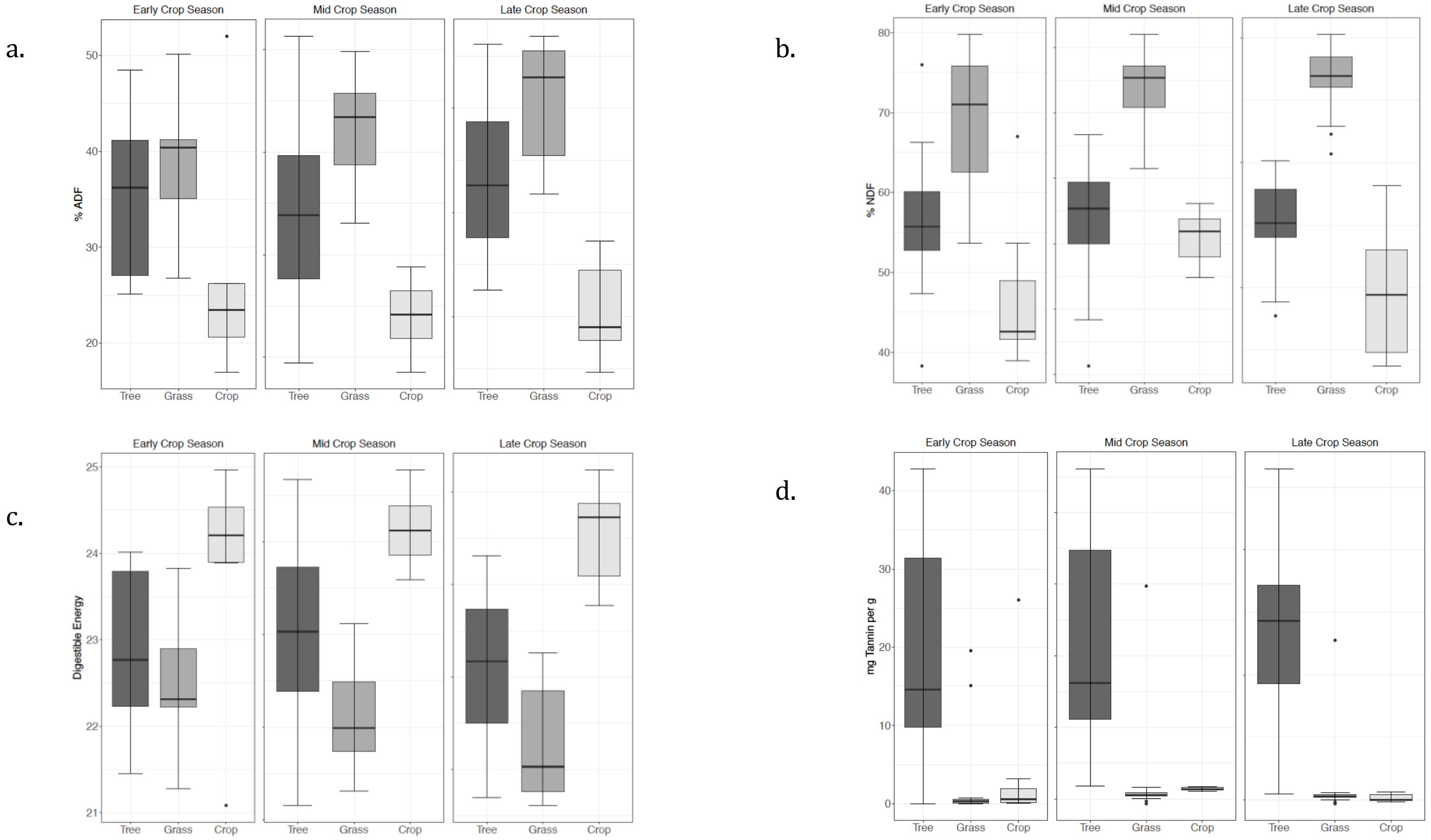


Figure 1. Boxplots comparing the differences in the vegetation characteristics a. ADF, b. NDF, c. Digestible Energy and d. Tannin between trees, grasses and crops, and their changes over the crop season.

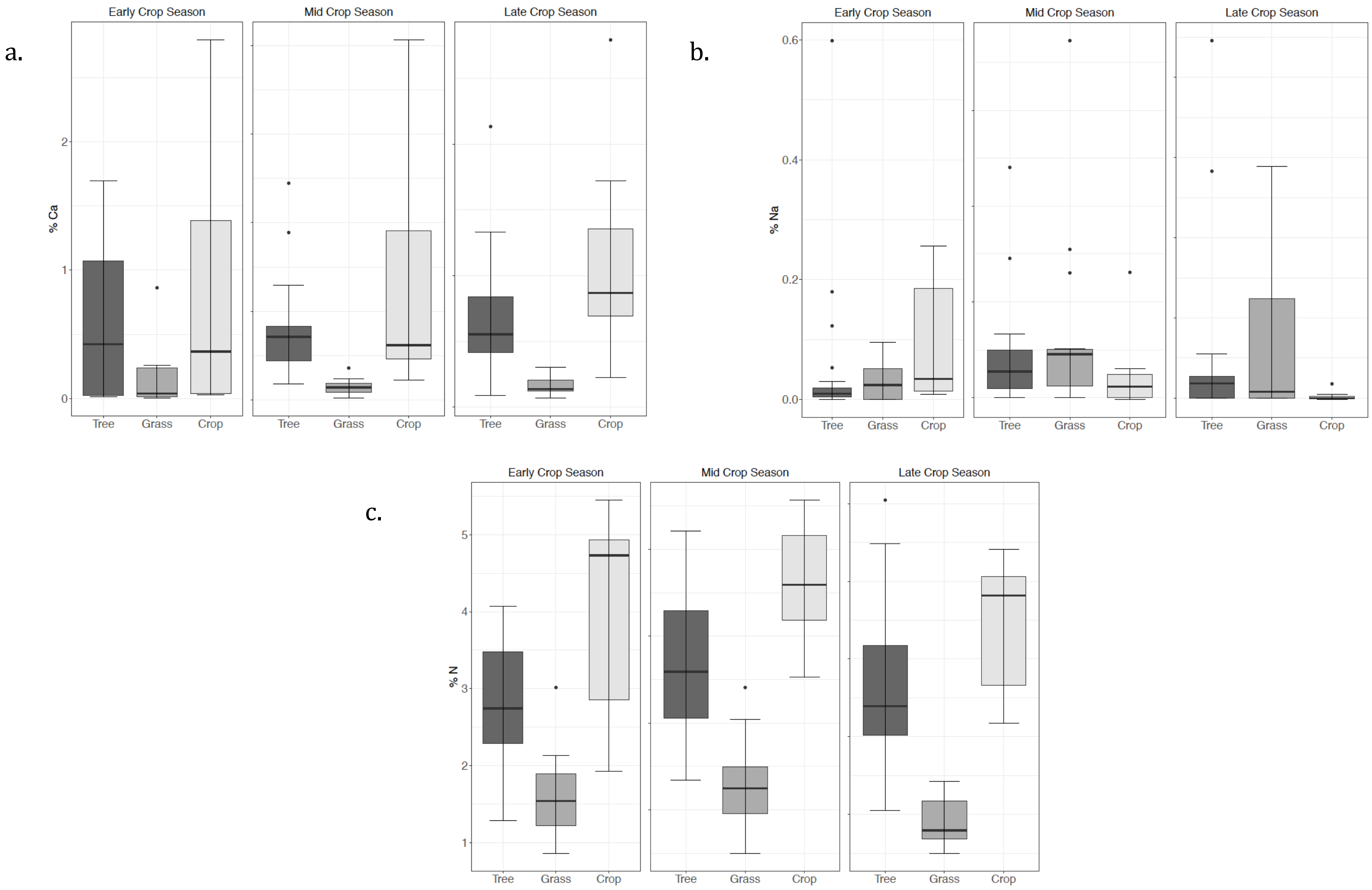


Figure 2. Boxplots comparing the differences in the vegetation characteristics a. calcium (Ca), b. sodium (Na), c. natrium (N) between trees, grasses and crops, and their changes over the crop season.

**Appendix VII**

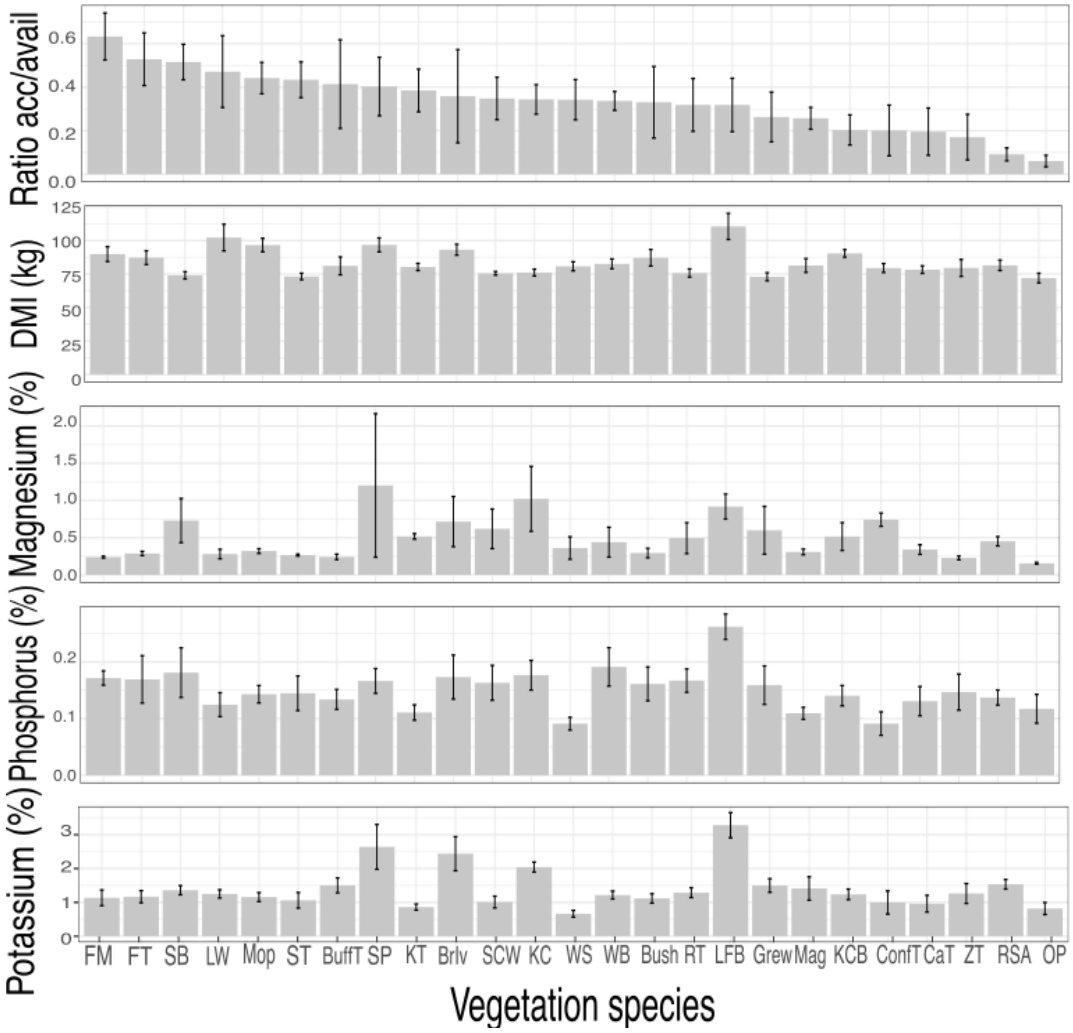


Figure 1. Plots of elephant dietary choices and their vegetation characteristics, sorted in order of acceptance/availability ratio.

**Appendix VIII – PCA results vegetation type comparisons and tree preference groups**

Table 1. PCA results comparing the vegetation characteristics of the three vegetation types tree, grass, crops early crop season.

| | **Comp. 1** | **Comp. 2** | **Comp. 3** |
|---|---|---|---|
| Eigenvalue | 5.799 | 1.131 | 0.825 |
| Percentage of Variance explained | 52.7% | 16.9% | 10.3% |
| Cum. Percentage of Variance explained | 52.7% | 69.7% | 80.0% |

| | **%NDF** | **%ADF** | **Tannin** | **%N** | **%P** | **%K** | **%Ca** | **%Mg** |
|---|---|---|---|---|---|---|---|---|
| Comp. 1 | -0.356 | -0.377 | | 0.355 | 0.350 | 0.327 | | 0.228 |
| Comp. 2 | 0.248 | | -0.224 | -0.125 | | 0.260 | -0.535 | 0.525 |
| Comp. 3 | 0.220 | -0.102 | -0.834 | -0.124 | | | 0.322 | -0.206 |

| | **%Na** | **DE** | **DMI** |
|---|---|---|---|
| Comp. 1 | 0.202 | 0.378 | 0.359 |
| Comp. 2 | 0.421 | | -0.231 |
| Comp. 3 | -0.143 | | -0.143 |

Table 2. PCA results comparing the vegetation characteristics of the three vegetation types tree, grass, crops, mid crop season.

| | **Comp. 1** | **Comp. 2** | **Comp. 3** |
|---|---|---|---|
| Eigenvalue | 5.365 | 1.998 | 1.214 |
| Percentage of Variance explained | 48.8% | 18.2 % | 11.0 % |
| Cum. Percentage of Variance explained | 48.8% | 66.9% | 78.0 % |

| | **%NDF** | **%ADF** | **Tannin** | **%N** | **%P** | **%K** | **%Ca** | **%Mg** |
|---|---|---|---|---|---|---|---|---|
| Comp. 1 | -0.360 | -0.404 | | 0.340 | 0.232 | 0.278 | 0.298 | 0.293 |
| Comp. 2 | 0.279 | | -0.598 | | 0.518 | 0.424 | | |
| Comp. 3 | -0.257 | -0.160 | | | 0.117 | | -0.577 | -0.594 |

| | **%Na** | **DE** | **DMI** |
|---|---|---|---|
| Comp. 1 | -0.104 | 0.405 | 0.332 |
| Comp. 2 | 0.209 | | -0.254 |
| Comp. 3 | 0.240 | 0.158 | -0.348 |

Table 3. PCA results comparing the vegetation characteristics of the three vegetation types tree, grass, crops late crop season.

| | **Comp. 1** | **Comp. 2** | **Comp. 3** |
|---|---|---|---|
| Eigenvalue | 5.925 | 1.598 | 0.833 |
| Percentage of Variance explained | 53.9% | 14.5% | 9.5% |
| Cum. Percentage of Variance explained | 53.9% | 68.4% | 78.0% |

| | **%NDF** | **%ADF** | **Tannin** | **%N** | **%P** | **%K** | **%Ca** | **%Mg** |
|---|---|---|---|---|---|---|---|---|
| Comp. 1 | -0.372 | -0.377 | | 0.316 | 0.326 | 0.187 | 0.316 | 0.301 |
| Comp. 2 | 0.205 | | -0.674 | -0.283 | 0.188 | 0.580 | | 0.133 |
| Comp. 3 | -0.158 | | 0.184 | -0.124 | -0.297 | | 0.221 | 0.241 |

| | **%Na** | **DE** | **DMI** |
|---|---|---|---|
| Comp. 1 | -0.118 | 0.377 | 0.363 |
| Comp. 2 | 0.158 | | |
| Comp. 3 | 0.849 | | |

*PCA trees and preference groups*

*Late Dry season*

The PCA of the tree characteristics across the year shows a different pattern than the PCAs comparing the three vegetation types. The first component exists out of all the characteristics except for calcium, with the fibre characteristics (NDF, ADF, DE and DMI) playing an important role. This component however only explains 40% of the variance and the other components all contain similar elements. Judging from the PCA biplots there is large scatter and overlap between the three preference groups in the late dry season (Table 7, Figure 8).

Table 4. PCA results on the vegetation characteristics of the three elephant dietary preference groups: preferred, intermediate and avoided in Late Dry season.

| | **Comp. 1** | **Comp. 2** | **Comp. 3** |
|---|---|---|---|
| Eigenvalue | 4.263 | 2.230 | 1.420 |
| Percentage of Variance explained | 38.8% | 20.3% | 12.9% |
| Cum. Percentage of Variance explained | 38.8% | 59.0% | 71.2% |

| | **%NDF** | **%ADF** | **Tannin** | **%N** | **%P** | **%K** | **%Ca** | **%Mg** |
|---|---|---|---|---|---|---|---|---|
| Comp. 1 | -0.428 | -0.398 | -0.178 | 0.343 | 0.283 | 0.166 | | 0.141 |
| Comp. 2 | -0.109 | -0.180 | -0.113 | -0.363 | -0.439 | | 0.522 | 0.406 |
| Comp. 3 | 0.297 | -0.371 | 0.177 | | -0.250 | -0.639 | | -0.420 |

| | %Na | DE | DMI |
|---|---|---|---|
| Comp. 1 | -0.220 | 0.399 | 0.427 |
| Comp. 2 | 0.368 | 0.180 | 0.109 |
| Comp. 3 | -0.186 | 0.369 | -0.288 |

*Early Wet season*

There are very small differences between the PCA of the late dry and the early wet season, with the components containing the same elements, yet the first component explains marginally more than that of the late dry season. The scatter plot however shows a different pattern that that of the late dry season, with a more concentrated low preference group positioned towards fibre and tannin (Table 8, Figure 8).

Table 5. PCA results on the vegetation characteristics of the three elephant dietary preference groups: preferred, intermediate and avoided in Early Wet season.

| | Comp. 1 | Comp. 2 | Comp. 3 |
|---|---|---|---|
| Eigenvalue | 4.263 | 2.230 | 1.420 |
| Percentage of Variance explained | 42.8% | 20.3% | 11.2% |
| Cum. Percentage of Variance explained | 42.8% | 63.1% | 74.4% |

| | %NDF | %ADF | Tannin | %N | %P | %K | %Ca | %Mg |
|---|---|---|---|---|---|---|---|---|

| | | | | | | | | | %Na | DE | DMI |
|---|---|---|---|---|---|---|---|---|---|---|---|
| Comp. 1 | -0.418 | -0.398 | -0.178 | 0.343 | 0.283 | 0.166 | | 0.141 | -0.220 | 0.399 | 0.427 |
| Comp. 2 | -0.109 | -0.180 | -0.113 | -0.363 | -0.439 | | -0.522 | 0.406 | 0.368 | 0.180 | 0.109 |
| Comp. 3 | | -0.371 | 0.177 | | -0.250 | -0.639 | | -0.420 | -0.186 | 0.369 | |

*Late Wet season*

Again, there is very limited difference with the PCA component information compared to the previous season, the early wet and late wet appear to have the same variance explanations. On the PCA biplot the low preference group appears to be concentrated even more, again around tannin and ADF and NDF (Table 9, Figure 8).

Table 6. PCA results on the vegetation characteristics of the three elephant dietary preference groups: preferred, intermediate and avoided in Late Wet season.

| | Comp. 1 | Comp. 2 | Comp. 3 |
|---|---|---|---|
| Eigenvalue | 4.263 | 2.230 | 1.420 |
| Percentage of Variance explained | 42.9% | 17.3% | 11.9% |

| | | | |
|---|---|---|---|
| Cum. Percentage of Variance explained | 42.9% | 60.2% | 60.2% |

| | %NDF | %ADF | Tannin | %N | %P | %K | %Ca | %Mg |
|---|---|---|---|---|---|---|---|---|
| Comp. 1 | -0.418 | -0.398 | -0.178 | 0.343 | 0.283 | 0.166 | | 0.141 |
| Comp. 2 | -0.109 | -0.180 | -0.113 | -0.363 | -0.439 | | 0.522 | 0.406 |
| Comp. 3 | | -0.371 | 0.177 | | -0.250 | -0.639 | | -0.420 |

| | %Na | DE | DMI |
|---|---|---|---|
| Comp. 1 | -0.220 | 0.399 | 0.427 |
| Comp. 2 | 0.368 | 0.180 | 0.109 |
| Comp. 3 | -0.186 | 0.369 | |

*Early Dry season*

During the early dry season there are again not many differences with the PCA component information compared to the previous season, however it is different from the late dry season when looking at the explaining of variance. The low preference group becomes less clustered and goes again to the more scattered pattern as in the late dry season biplot (Table 10, Figure 8).

Table 7. PCA results on the vegetation characteristics of the three elephant dietary preference groups: preferred, intermediate and avoided in Early Dry season.

| | Comp. 1 | Comp. 2 | Comp. 3 |
|---|---|---|---|
| Eigenvalue | 4.263 | 2.230 | 1.420 |
| Percentage of Variance explained | 43.1% | 16.1% | 13.7% |
| Cum. Percentage of Variance explained | 43.1% | 59.3% | 73.0% |

| | %NDF | %ADF | Tannin | %N | %P | %K | %Ca | %Mg |
|---|---|---|---|---|---|---|---|---|
| Comp. 1 | -0.418 | -0.398 | -0.178 | 0.343 | 0.283 | 0.166 | | 0.141 |
| Comp. 2 | -0.109 | -0.190 | -0.113 | -0.363 | -0.439 | | -0.522 | 0.406 |
| Comp. 3 | | -0.371 | 0.177 | | -0.250 | -0.639 | | -0.420 |

| | %Na | DE | DMI |
|---|---|---|---|
| Comp. 1 | -0.220 | 0.399 | 0.427 |
| Comp. 2 | 0.368 | 0.180 | 0.109 |
| Comp. 3 | -0.186 | 0.369 | |

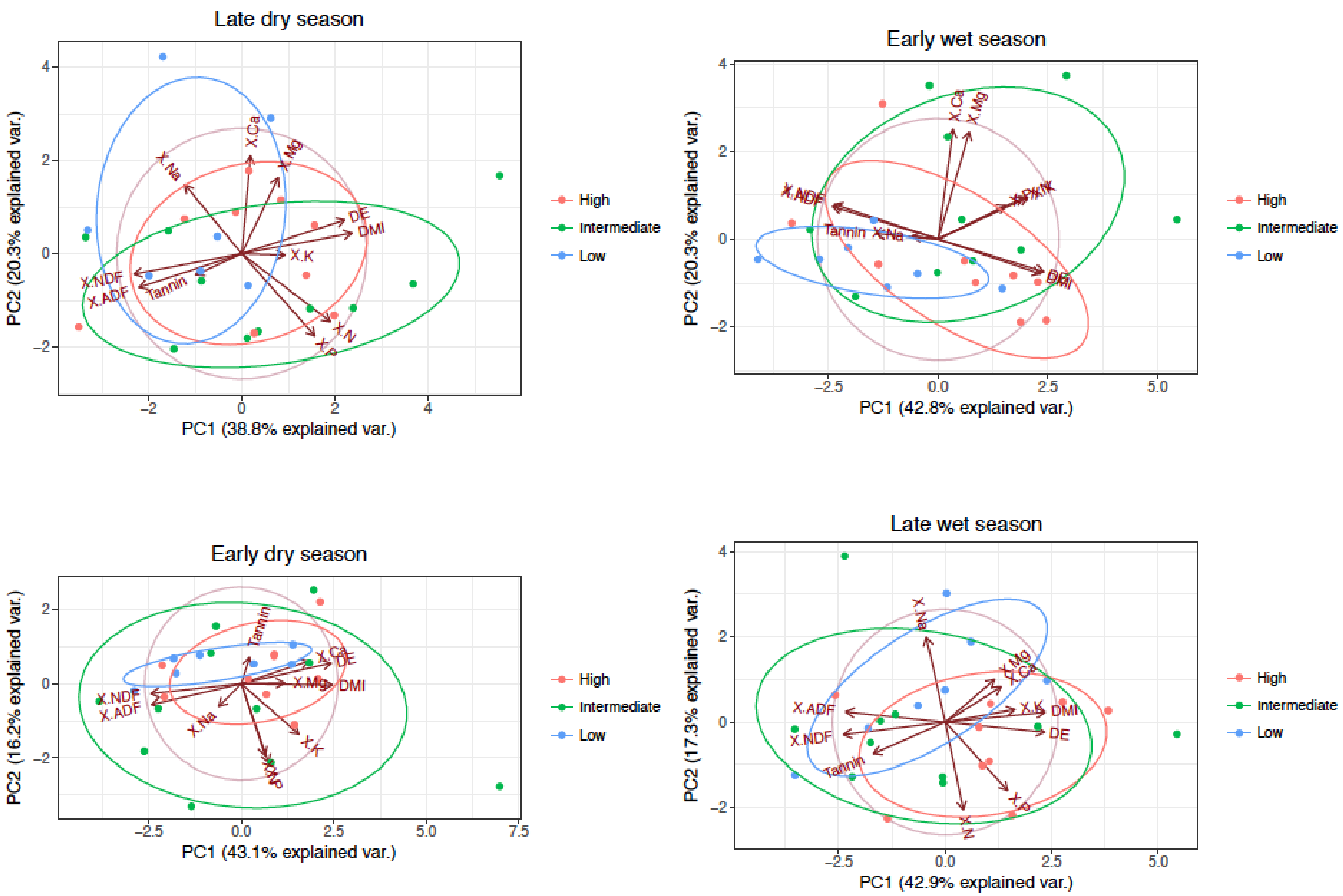


Figure 1. Biplots of PCAs for tree characteristics the four seasons, revealing the clusters of elephant preference group.

### Appendix IX - Nutritional Geometry

Nutritional Geometry methods, in particular Right-angle Mixture Triangles (RMTs) plot the ideal nutrient balance for animals, and the nutrient balance of different food sources available to them, in order to analyse how animals can reach their nutritional requirements by combining the food sources (Raubenheimer and Simpson, 1993, 1999; Simpson *et al.*, 2004). RMTs are three dimensional spaces plotted on a two dimensional surface, showing the percentages in which different components are present in a composition, demonstrating nutrient balances and ideal compositions (figure 9.1; Raubenheimer, 2011).

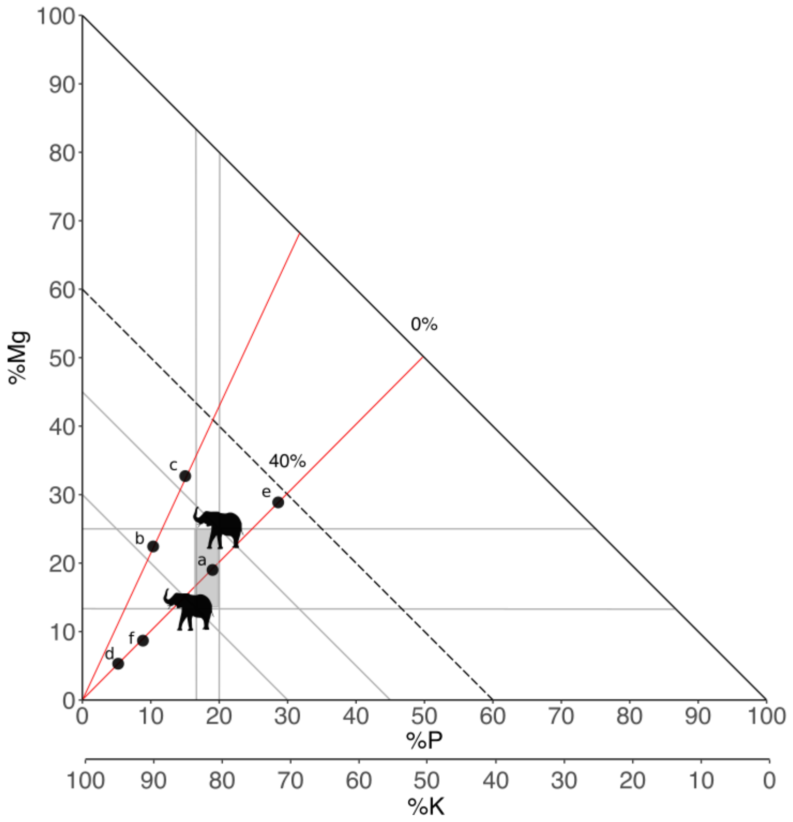

*Figure 3.1 Right-Angle Mixture Triangle.*

RMTs do not reflect actual nutritional requirements, but demonstrates how balanced the food items are in their micronutrient composition, and how the elephant could combine food items to achieve the balanced diet the elephant requires. In figure 3.1, the X-axis represents phosphorus (P) %, the Y-axis magnesium (Mg) % and the diagonal Z-axis potassium (K) %. The grey square between the two elephant icons indicates the nutrient space. The position of the elephants is based on the upper and lower limits of the required dietary balance of Mg:P:K for elephants. The third axes starts at the base of the triangle, reaching 40% at the dotted line. For each element the two grey lines derived from the elephant indicate the nutrient space in which that individual element is balanced. We plotted 6 hypothetical food items, with food *a* in the nutrient space of the required Mg:P:K balance. Food item *b* has the correct Mg:K balance, but falls short on the percentage of P (10%), while item *c* shows a deficiency in both P (15%) and K (53%), and a higher density of Mg than required by the elephant (32%). Food item *d* has an excessively high K-percentage of 90% while it is deficient in both P and Mg. Combinations of food items can be complementary if they are aligned on the red lines from the origin of the plot through the required dietary points, but fall on opposite sides of the intake target, as food items *d* and *e*, or are substitutable if they fall on the same side of the intake target, which is the case for *d* and *f*, which both could complement *e* (Raubenheimer, 2011).

http://www.jstor.org/stable/3543921.
